# Dictionary of human intestinal organoid responses to secreted niche factors at single cell resolution

**DOI:** 10.1101/2025.05.06.652311

**Authors:** Meghan M. Capeling, Bob Chen, Kazeera Aliar, Elisa Penna, Veronica Ibarra Lopez, Conrad Foo, Sandra Rost, Loryn Holokai, Xinming Tong, Devan Phillips, Caden Sweet, Jing Li, Sharmila Chatterjee, Elizabeth Skippington, Zora Modrusan, Lisa M. McGinnis, Runmin Wei, Mary Keir, Orit Rozenblatt-Rosen, Michelle B. Chen

## Abstract

The intestinal epithelium is often a site of pathology, such as in inflammatory bowel disease (IBD), and its maintenance is highly modulated by interactions with the microenvironment. However, a systematic understanding of how the myriad of niche cues impact distinct epithelial cell types in a diseased context is still lacking. To address this gap, we first benchmarked diverse human colonic organoid injury models against IBD tissue, and established a disease-relevant model of epithelial inflammation using TNFα, IFNγ, and IL1β. Using this system, we built a dictionary of epithelial responses to 81 secreted niche factors at single cell resolution via donor-pooled, multiplexed single cell RNA-sequencing (scRNA-seq). The comprehensive nature of our atlas allowed us to map relationships between perturbations, infer the function of less well-characterized ligands, and identify cell type-specific perturbed pathways. Finally, we established the relevance of organoid-derived gene programs by mapping them to single cell and spatial atlases of human IBD tissue. Our resource offers a global view of epithelial responses to microenvironmental cues in a physiologically relevant disease context and generates new hypotheses for signaling factors that may be involved in epithelial homeostasis and repair.

## INTRODUCTION

The colonic epithelium is a highly regenerative tissue composed of proliferative stem cell-containing crypt domains and differentiated cell types that facilitate absorption and mucus secretion^1^. During homeostasis, spatially-patterned microenvironmental cues direct stem cell proliferation in the crypts, which supports turnover and differentiation into specialized cell types in the inter-crypt domains^2^. However, the microenvironment becomes altered in the context of injury or disease, particularly inflammatory bowel disease (IBD), a common intestinal disorder characterized by chronic inflammation^3^. For instance, stromal populations in the stem cell niche are altered during IBD, which may lead to disrupted signaling to the epithelium ^4,56^.

While much is known about the colonic microenvironment during homeostasis and development^1,2,7^, it is less clear how niche cues contribute to epithelial repair in a diseased context. First, epithelial regeneration is most commonly studied in models which lack important aspects of disease biology, including mouse models which suffer from species-specific differences,^8,9^ immortalized cell lines or 2D cultures which do not capture the complex 3D architecture of the gut^10^, or healthy homeostatic organoids that do not model regeneration in the face of injury. Organoids derived from IBD patients have been employed to overcome these issues^11^, but they often lose their inflammatory phenotype over time in culture^12^. Second, the lack of systematic interrogation of epithelial cell-type specific responses to niche cues hinders a global understanding of pathway regulation associated with homeostasis and repair. Most studies to date have focused on characterizing a select few perturbations at a time,^7,13,14^ rendering it difficult to map out divergences and similarities between ligands. Conversely, large phenotypic screens without molecular profiling are also powerful but pose challenges in gaining direct mechanistic understanding^15^. Moreover, because the intestinal epithelium is heterogenous in nature, understanding cell-type specific responses at single cell resolution can provide insight on both mechanism and therapeutic targeting. Indeed, recent scRNA-seq chemical perturbation screens have already begun to demonstrate value in unbiased and systematic perturbation mapping^16,17,18^.

To address these gaps, we first developed a physiologically relevant epithelial injury model by characterizing the response of healthy human colon organoids subjected to common epithelial stressors (gamma irradiation, cytokine stimulation, dextran sodium sulfate (DSS), and mechanical injury)^19,20,21,22,23,24^ and benchmarking them to human IBD tissue. We discovered an IBD-relevant inflammatory model and used it to systematically profile epithelial responses to 81 secreted niche factors known or predicted to interact with the human colonic epithelium during development, homeostasis, and disease^25–27^ via scRNA-seq. The breadth of our atlas allowed us to infer cell type specific effects of less well-characterized ligands via their relationships to known ligand classes, and identify epithelial response gene programs that may be implicated in a return to homeostasis. Overall, we present a valuable resource to probe the diversity and context-specificity of human intestinal epithelial responses to endogenous microenvironmental cues, which can be leveraged for therapeutic target discovery and biomarker selection for IBD.

## RESULTS

### Development of a human colonic organoid injury model that recapitulates molecular features of human IBD

We developed 4 epithelial injury models starting from homeostatic primary colon organoids derived from 3 human donors. Organoids were cultured in a commercial microwell suspension culture system^28^ to improve ease of imaging, single-cell dissociation, and homogeneity. We confirmed that microwell-grown organoids contain both stem and differentiated cell types, and display transcriptional similarity to both traditional organoids cultured in 3D extracellular matrix domes and to fresh human colon tissue (Extended Data Fig. 1, Table S1). Organoids were first differentiated to mimic the complexity of cell types found *in vivo* and then treated with conventional epithelial stressors including DSS, gamma irradiation, and cytokines (TNFα, IFNγ, IL1β combined - termed cytomix) ^24,29^ across various dosages **(Fig. 1A)**. We first characterized organoid responses via morphology, degree of cell death, and barrier function^30^, as IBD lesions display increased expression of apoptotic markers^31^ and decreased expression of tight junction proteins leading to increased permeability^32^. DSS and cytomix treatment resulted in morphological changes from bright and cystic to dark organoid structures, suggesting stress and cell death **(Fig. 1B)**. While control and irradiated organoids were primarily simple squamous or columnar in architecture, DSS-treated organoids showed epithelial thickening and altered polarity, with Phalloidin staining both the apical and basolateral surfaces. In contrast, cytomix treatment resulted in primarily pseudostratified epithelium, which has been reported for IBD patient-derived organoids^33^. Out of all treatments, only DSS caused significant decreases in proliferation (**Fig. 1B**, Extended Data Fig. 2A). While irradiation has previously been shown to decrease proliferation in small intestinal organoids^21,14^, we did not observe decreases in KI67 expression at 6Gy or 12Gy, indicating potential differences in susceptibility between the small and large intestine.

**Figure 1:**
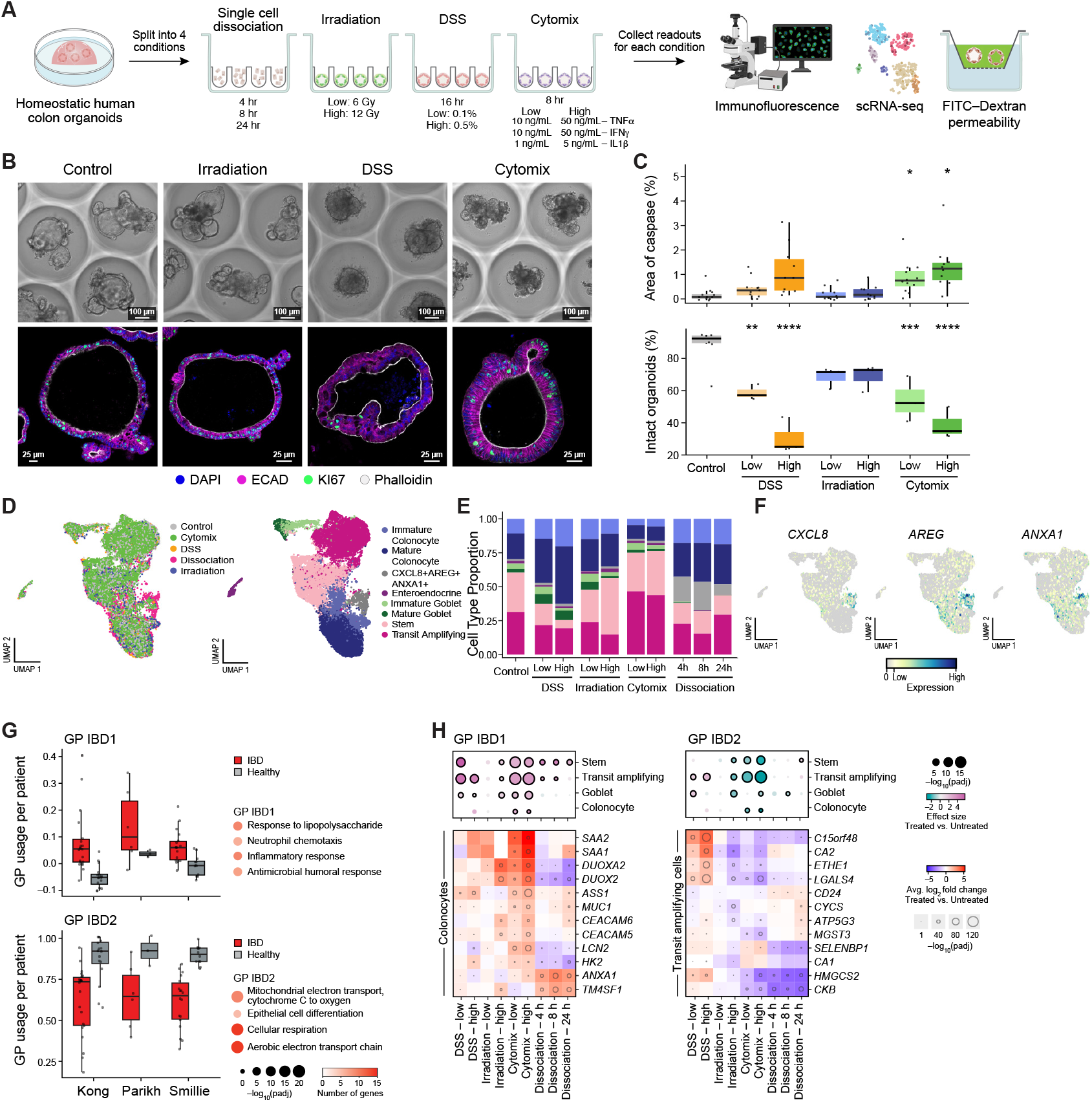
Development of a human colonic organoid disease model that recapitulates human IBD molecular phenotypes. **(A)** Development and characterization of four injury models: organoids were either dissociated and allowed to recover from single cells, γ-irradiated at 6Gy or 12Gy, treated with 0.1% or 0.5% DSS for 16 hours, or treated with cytomix (combination of IFNγ, TNFα, IL1b) for 16 hours. **(B)** Representative brightfield and IF images (ECAD, Phalloidin, KI67, DAPI) of injury conditions at one day post-injury. Scale bar = 100μm (brightfield), 25μm (IF). **(C)** Quantification of apoptosis (top) and permeability (bottom) of organoids 1 day post-injury, n>=10 organoids per condition, 3 donors tested. **(D)** Dimensional reduction of all combined injured organoids, collected one day-post injury for DSS and cytomix-treated organoids, or 8, 16, and 24 hours post-dissociation. **(E)** Cell type composition changes across injury conditions. **(F)** Expression of *AREG, CXCL8*, and *ANXA1*, enriched in the dissociation-enriched cluster. **(G)** Left: Average module usage of GPs IBD1 and IBD2, per patient, in 3 published IBD atlases. Right: Top enriched gene ontology (GO) terms for modules IBD1 and IBD2. **(H)** Top: Effect size (treated/non-treated) of module IBD1 (left) and module IBD2 (right) across cell types and injury conditions. Bottom: LogFC of genes in treated vs. untreated organoids, for colonocytes (IBD1) and transit amplifying cells (IBD2).

Finally, activated Caspase-3/7 staining revealed dose-dependent increases in apoptosis after DSS and cytomix, but not irradiation (**Fig. 1C**, Extended Data Fig. 2B, Table S2)^34^. All injuries except for irradiation led to a significant increase in organoid permeability to 4kDa FITC dextran compared to untreated organoids, with strong dose-dependent effects in both DSS and cytomix treatment (**Fig. 1C**, Extended Data Fig. 2B, Table S2). These results suggest that DSS and cytomix elicit a functional stress response in human colon organoids that may bear functional relevance to human IBD.

To understand the molecular fidelity of these injury models to human IBD, we performed scRNA-seq after one day of treatment (**Fig. 1D**). Changes in cell type proportions were observed, including decreased stem and transit-amplifying (TA) cells with DSS, and decreased colonocytes and increased TA/stem cells with cytomix (**Fig. 1E**). Additionally, a unique cell cluster marked by expression of *CXCL8, AREG*, and *ANXA1* was enriched specifically in organoids recovering from dissociation (**Fig. 1F**). This population may represent a unique cell state brought on by endogenous organoid formation mechanisms, as *AREG* and *ANXA1* have been reported as markers of regeneration and de-differentiation in organoids and mouse models.^19,35,36^ To determine the disease-relevance of organoid molecular responses, we first conducted a meta-analysis across two published single-cell datasets (colon tissue from healthy, Crohn’s disease, and ulcerative colitis patients) to identify conserved gene programs (GPs) dysregulated between diseased vs. healthy epithelium ^4,25^ (Extended Data Fig 2C, Table S3). Two highly conserved gene programs were found: upregulation of GP ‘IBD1’, which is associated with inflammation (*DUOX2, DUOXA2, CEACAM6, ANXA1)*^*37*^ and contains known IBD risk genes^38,39^, and downregulation of GP ‘IBD2’, which is associated with cellular differentiation (*CD24, CA1, KRT19)* and metabolism (*COX8A, NDUFB9, CYCS*), particularly oxidative phosphorylation^4^ and electron transport (**Fig. 1G**). These changes occur across most epithelial cell types (Extended Data Fig. 2C), suggesting a general heightened immune response and altered metabolic function, as previously described.^40–45^ Additionally, IBD1 and IBD2 appear highly predictive of disease state (Extended Data Fig. 2D) and show similar trends when tested in a third unseen dataset^46^ (**Fig. 1G**). We then compared the expression of IBD1 and IBD2 across injury models. IBD1 was upregulated in stem and TA cells under DSS, and even more strongly with cytomix in a dose-dependent manner (**Fig. 1H**, Table S4, Table S12). Conversely, metabolism-related IBD2 was only downregulated in multiple cell types after cytomix treatment, and to a lesser extent with irradiation. Combined, these results suggest that cytomix stimulation bears the strongest phenotypic and molecular resemblance to human IBD tissue.

We additionally identified GPs associated with each injury in an unbiased manner (**Fig. 2A**, Extended Data Fig. 3A,B, Table S5, Table S12). Dissociation upregulated wound healing and apoptosis pathways (GP19: *DKK1, ANXA1, ANXA5*), processes associated with regeneration. Cytomix upregulated immune response and interferon pathways (GP16: *DUOX2, DUOXA2, CXCL10*), irradiation altered DNA damage, apoptosis, and translation pathways (GP8: *LY6D, BAX, BCL2L1*), while DSS upregulated transcription and cytoskeleton reorganization processes (GP6: *THRB, HNF4G, LGR4*). We additionally identified a general organoid injury signature (GP2: *LDHA, ME1, ENO1*) which was downregulated across all injuries and associated with metabolic and biosynthetic processes, and may indicate a broad metabolic defect across injured epithelium (Extended Data Fig. 3C, Table S5).

**Figure 2:**
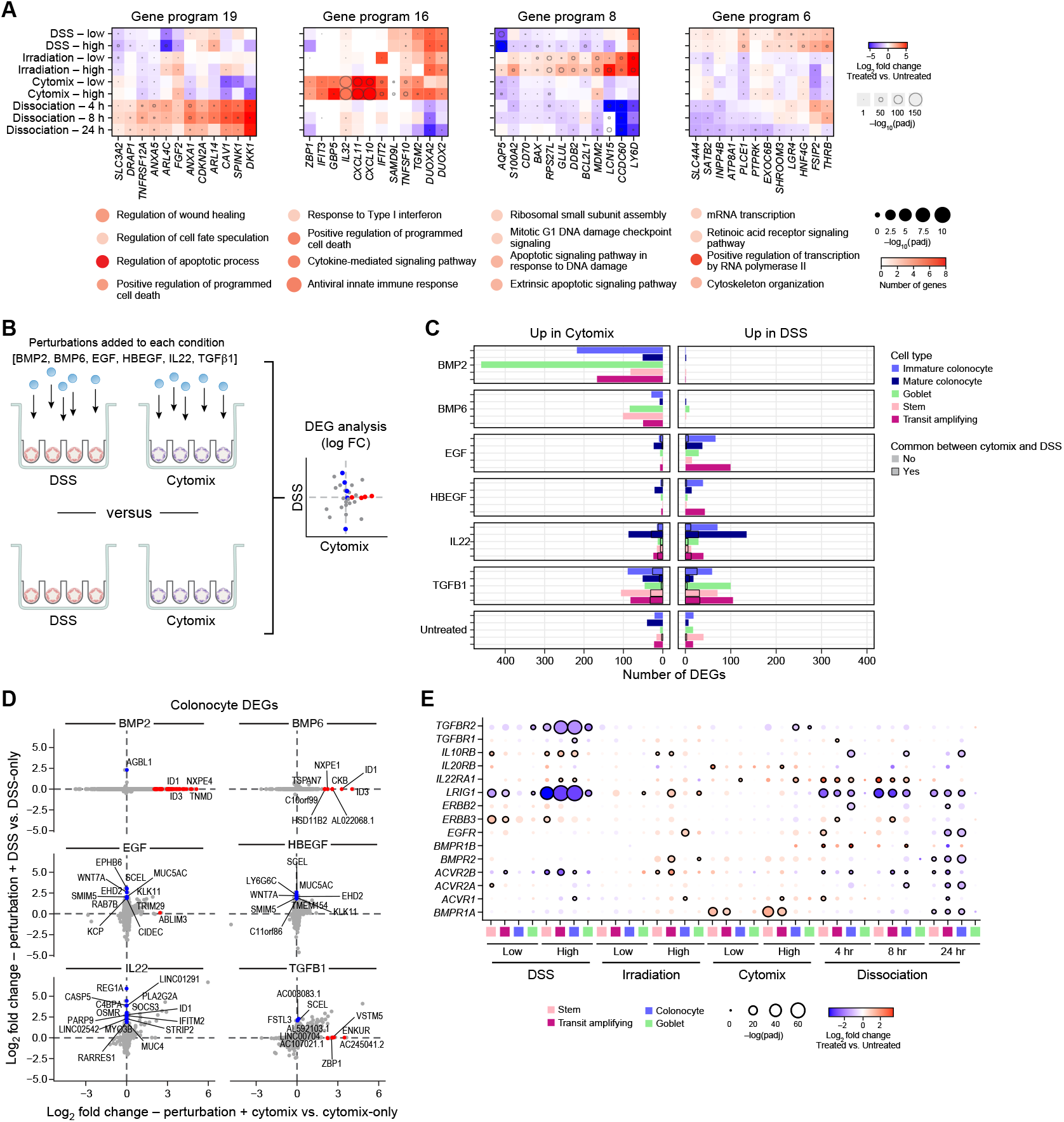
Epithelial responses to perturbations are context-specific to starting injury states. **(A)** Top: LogFC (injury/no injury) of top genes from select cNMF-derived gene programs across injury types (DSS, irradiation, cytomix, dissociation) and dosages, in TA cells. Bottom: Top GO terms associated with each gene program, where circle size represents -log(padj) and circle color represents number of genes per GO term. **(B)** Overview of experiment testing the effect of the same perturbations (EGF, HBEGF, BMP2, BMP6, TGFβ1, IL22) on different injury model backgrounds (DSS vs cytomix). **(C)** Number of upregulated DEGs (padj <0.05, logFC > 1) for each perturbation when applied to either the DSS model (right) or cytomix model (left). Colors depict distinct epithelial cell types. Black outlines depict the number of DEGs that are shared between both models upon treatment. **(D)** Comparison of log2-fold change of DEGs in colonocytes for each secreted factor applied to either the cytomix model (x axis) or the DSS model (y axis) (red=exclusively DE in cytomix-treated organoids, blue=exclusively DE in DSS-treated organoids). **(E)** Average log2-fold change of receptors and antagonists for applied perturbations in each injury model (injured/ non-injured)

**Figure 3.**
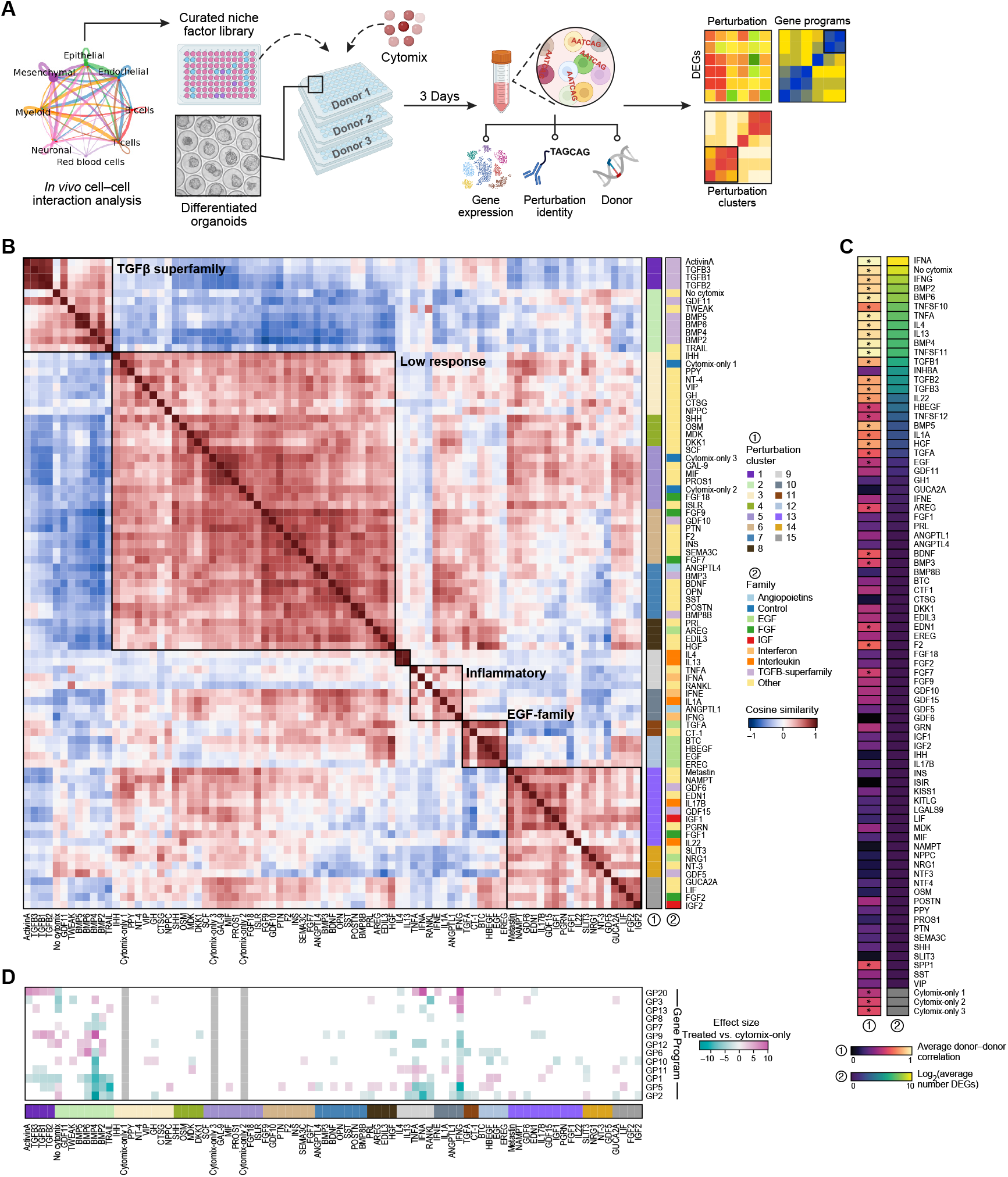
Landscape of colonic epithelial cell responses to secreted niche factors in an inflamed background at single cell resolution. **(A)** Overview of experimental design. High probability secreted factors signaling between the intestinal niche and the epithelium were computationally inferred from a meta-analysis of 3 human datasets. Human recombinant proteins corresponding to 81 secreted ligands were applied to inflamed differentiated intestinal organoids in an arrayed format in microwell plates across 3 donors. Donors were pooled and perturbations were hashed using HTOs and subject to scRNA-seq and perturbation/donor demultiplexing. **(B)** Left: Cosine similarity of all perturbations (3 donors averaged) grouped by perturbation clusters. Perturbations are annotated by perturbation ligand family class and perturbation cluster (determined by k means clustering (k=12)). **(C)** Left: Reproducibility of response across donors represented by the average cosine similarity between 3 donors within a given perturbation. Asterisks indicate cosine similarity is higher than 95% percentile of permuted distribution (rho=0.406). Right: Average number of differentially expressed genes between each perturbation vs cytomix-only control across each major cell type per perturbation (n=3 donors per comparison). **(D)** Effect size (perturbation/no perturbation) of activity GPs (all cell types averaged) across 3 donors. Only p.adjusted<0.05 are colored.

To assess the IBD-relevance of injury-elicited GPs, we evaluated their expression in human IBD tissue. Only cytomix-associated GPs (GP3, GP16) exhibited concordant upregulation in two human IBD datasets^4,25^ (Extended Data Fig. 4A, Table S11). We additionally examined the expression and localization of these GPs in healthy (n=4) vs UC (n=16) patient samples via spatial transcriptomics (CosMx, Extended Data Fig. 4B, Table S10A). Importantly, cytomix-regulated GP3 was enriched in epithelial cells in UC vs healthy, while most other stressor induced GPs did not exhibit enrichment in disease (Extended Data Fig. 4C).

**Figure 4:**
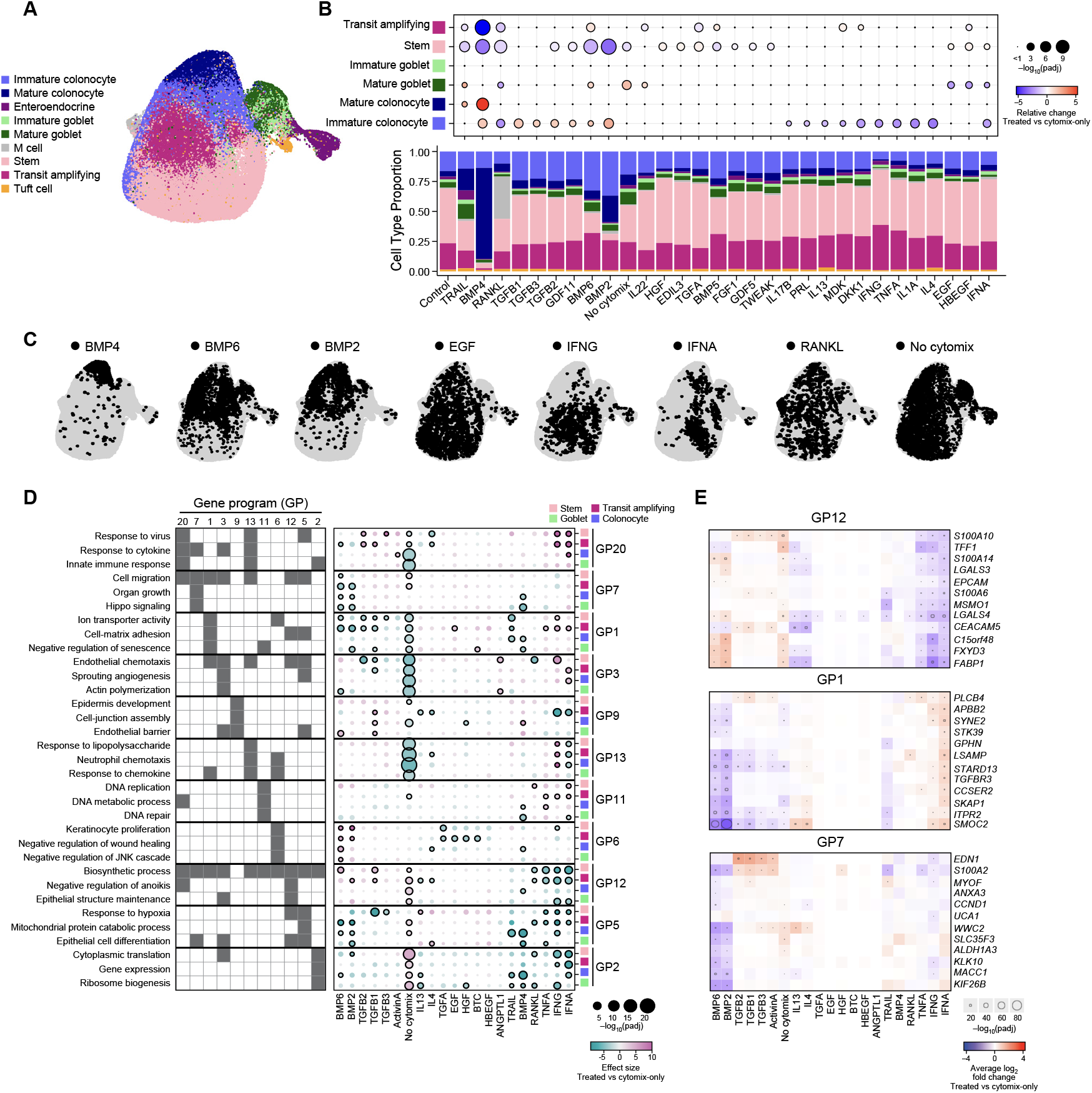
Niche factor perturbations elicit distinct transcriptional programs that are cell type specific. **(A)** Dimensional reduction of combined perturbation landscape annotated by epithelial cell types. **(B)** Proportion of cell types per perturbation (barplot) and the magnitude and direction of the change in proportion (coefficient) between treated vs cytomix-only organoids (n=3 donors). **(C)** Location of specific perturbations of interest in UMAP embedding. **(D)** Left: Top GO terms for select activity GPs across a select list of high impact perturbations. Right: Effect size of select activity GPs across perturbations in distinct cell types (transit amplifying, stem, goblet, colonocyte). Statistically significant effect sizes (padj < 0.05) are outlined in black. **(E)** Log2FC (treated/cytomix-only)) of top genes for GPs 1, 7, 12 and select perturbations in transit amplifying cells.

Together, this suggests that cytomix treatment best recapitulates IBD-like epithelial gene signatures, while conventional stressors (DSS, irradiation) bear less relevance when applied to human colon organoids.

### Epithelial responses to perturbations are context-specific based on starting injury states

We further hypothesized that organoid responses to perturbations may be primed by stressors and hence be context-specific. To test this, we performed scRNA-seq on DSS and cytomix treated-organoids perturbed with growth factors with known roles in intestinal epithelial cell maintenance: BMP2, BMP6, EGF, HBEGF, TGFβ1, and IL22 (**Fig. 2B**) ^13,14,47,48^. Notably, we found significantly more differentially expressed genes (DEGs, padj < 0.05, logFC > 1) for BMP6 and BMP2 in cytomix-treated organoids compared to DSS, while DSS-treated organoids exhibited many more DEGs in the presence of EGF and HBEGF (**Fig. 2C**, Extended Data Fig. 5A,B, Table S6). BMP2/6 strongly upregulated canonical BMP target genes (*ID1, ID3*) and genes involved in steroid (*GUCY2C, AKR1C3*) and sphingosine (*ACER2, ACER3*) metabolism and epithelial maintenance and mucus production (*MUC13, TFF1, MUC4*) specifically in the context of cytomix, while EGF ligands elicited pathways involving regulation of cell-cell adhesion and anoikis (*CEACAM5, CEACAM6, CAV1*) and cell-substrate adhesion (*COL17A1, ITGB4, RHOD*) specifically in DSS (**Fig. 2D**, Extended Data Fig. 4A-C). We hypothesized that these differences could be partially attributed to changes in expression of receptors and inhibitors of both pathways after injury. For example, cytomix treatment upregulated BMP receptor *BMPR1A* while DSS upregulated EGF-family receptors *ERBB2* and *ERBB3* and downregulated EGFR-inhibitor *LRIG1* (**Fig. 2E**). Interestingly BMP receptors (*BMPR1A, ACVR1*) and *EGFR* are upregulated in IBD vs. healthy controls while *ERBB2* is downregulated (Extended Data Fig. 5D). Additionally, response genes may already be activated by the injury itself as cytomix-treated organoids upregulated IL22 response genes while DSS upregulated BMP2 response genes (Extended Data Fig. 5E), thus decreasing the dynamic range to observe perturbation responses. This suggests that the choice of injury model plays an important role in evaluating physiologically relevant epithelial responses, and further strengthens the selection of cytomix as an IBD-relevant model^49^.

**Figure 5:**
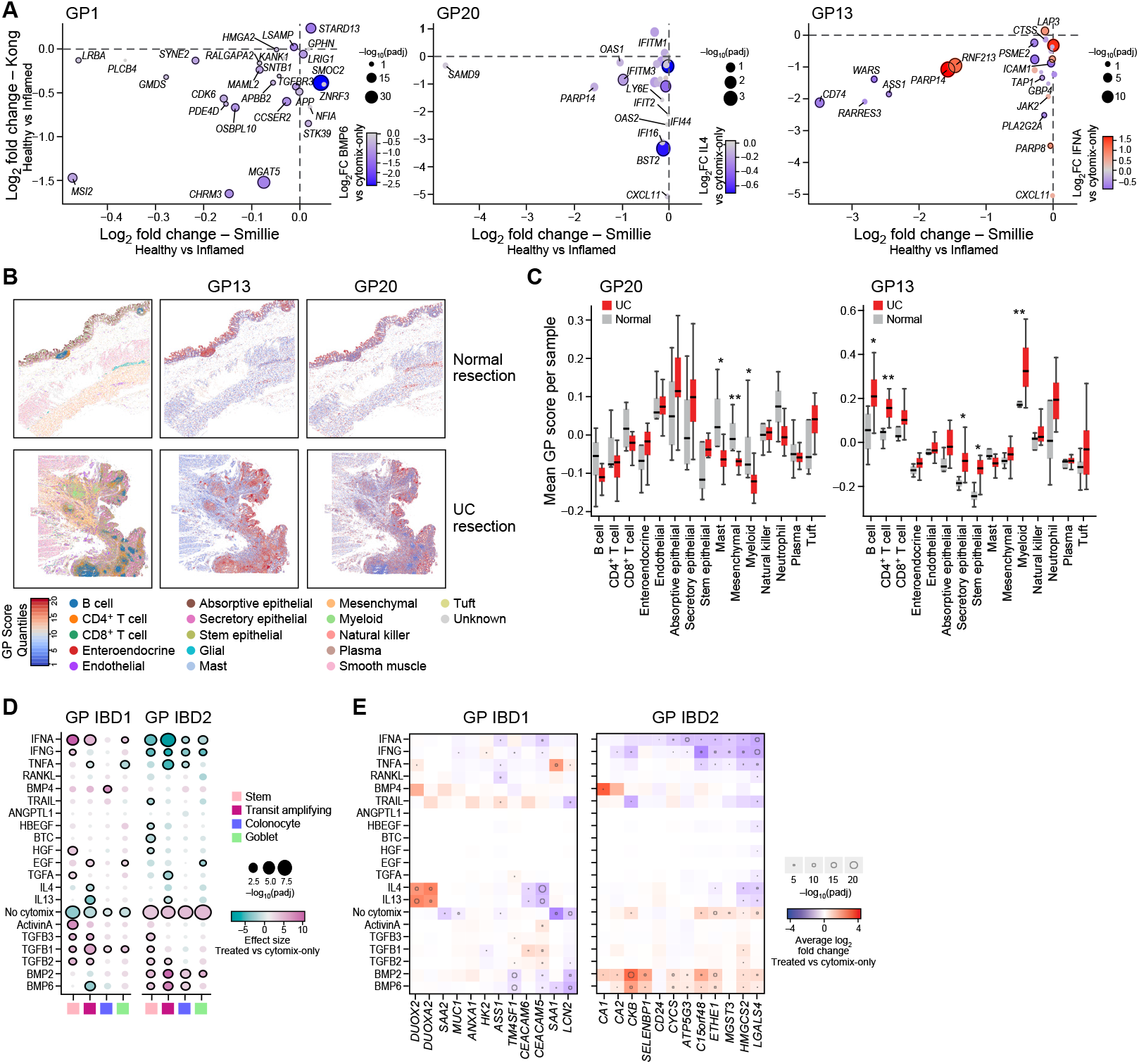
Perturbation-elicited organoid gene programs are active in human IBD tissue. **(A)** Log2FC (healthy vs. inflamed) of GP1, GP3, and GP20 genes in the epithelium of human IBD datasets^4,25^. Dots are sized by log2FC in perturbation treated organoids (BMP6, IFNα, IL4) vs. cytomix-only organoids, with size corresponding to -log10(padj). All log2FC values represent transit amplifying cells. Black outlines indicate padj <0.05. **(B)** Representative CosMx images of healthy vs. UC tissue, highlighting cell type annotations and expression scores of GP13 and GP20. **(C)** Mean expression of GP13 and GP20 per sample in UC vs. healthy tissue (right), *p<0.05, **p<0.01, and ***p < 0.001. **(D)** Effect size of IBD tissue-derived GPs IBD1 (inflammation) and IBD2 (metabolism) in select perturbations. **(E)** Log2FC (treated/cytomix-only) of top IBD1 and IBD2 genes in organoids. Black outlines indicate padj <0.05.

### Landscape of colonic epithelial cell responses to secreted niche factors in an inflamed background at single cell resolution

Using the low-dose cytomix model, we generated a molecular dictionary of epithelial responses to secreted niche factors in human colon organoids across 3 donors via multiplexed scRNA-seq (**Fig. 3A**). We selected a perturbation space enriched in factors predicted to promote epithelial homeostasis, wound healing, and/or regeneration by performing a meta-analysis of putative ligand-receptor interactions between the gut epithelium and its niche via existing single cell atlases (Extended Data Fig. 6A, Table S7).^27,25,26^ We selected 70 niche factors from this analysis and 9 additional perturbations with known effects on the intestinal epithelium such as IL22^48^, DKK1^50^, and ISLR^51^, plus 2 controls, for a total of 81 perturbations (36.8% predicted to come from stromal, 23.2% immune, 16.8% neuronal, and 23.2% epithelial). Recombinant human proteins (Table S14) were applied to differentiated cytomix-treated colon organoids from 3 donors in an arrayed format followed by multiplexed scRNA-seq to assess transcriptional responses **(Fig. 3A)**. After filtering and quality control, we obtained 169,996 cells with 3 donors x 81 perturbations, for a total of 243 independent conditions (Extended Data Fig. 6B).

To map the overall landscape of relationships between perturbations, we calculated the cosine similarity between all perturbations (**Fig. 3B**). Ligands in clusters 3-8 (‘Low Response’) display similarities to cytomix-only controls, suggesting that the majority of these perturbations did not elicit an identifiable response over the injury itself. Perturbations from the same family generally clustered together with high similarity at the pseudo-bulked level (EGF-family ligands, TGFβ-superfamily ligands, pro-inflammatory molecules IFNε, TNFα, IFNγ), demonstrating the ability to recover known ligand pathway redundancies. We also uncovered divergences in activity across ligand families, including NRG1 clustering with SLIT3 and NT-3 instead of other EGF-family ligands, potentially due to EGF-like domains within SLIT3, and TRAIL(*TNFSF10*)/TWEAK(*TNFSF12*) clustering with the TGFβ-superfamily rather than RANKL(*TNFSF11*) and TNFα, suggesting disparate responses in the context of the inflamed epithelium. Interestingly, uninjured (no cytomix) organoids clustered with members of the TGFβ-superfamily, including BMP6 and GFD11, suggesting that BMP treatment on inflamed organoids may elicit a transcriptional response that drives higher similarity to non-inflamed organoids. Highly correlated perturbations are also preserved at the cell type specific level, indicating that similarities are largely not driven by cell type composition differences.

To determine the genes driving these relationships, we performed DEG analysis for every perturbation vs. cytomix-only controls in a cell-type specific manner (Extended Data Fig. 7A-D, Table S8). 25/80 perturbations elicited a response with more than 1 DEG (false discovery rate (FDR) < 0.05, **Fig 3C**). In particular, members of the BMP, TGFβ, TNF, and IL families exhibited strong transcriptional responses with high numbers of DEGs (>25 based on p-adj < 0.05, abs(log2FC) >1) and donor reproducibility (**Fig 3C**).

We recovered known biology as ligands upregulated canonical target genes, including BMP2/6 (*ID1, ID3*), TGFβ1/2/3 (*SMAD3, CDKN1A*), and IL22 (*DMBT1, MUC1*)^49^. We additionally uncovered less well-known responses which may be specific to an inflamed context, including BMP2/6 upregulating genes associated with GP IBD2 (*CKB, SELENBP1, C15orf48*) suggesting a role in maintaining metabolic homeostasis, TGFβ1/2/3 upregulating interferon response genes (*SP100, CX3CL1*) implicating their roles in epithelial response to inflammation, and TRAIL downregulating pro-apoptotic genes (*BNIP3L, FAM162A, BNIP3*) suggesting an anti-inflammatory role (Extended Data Fig. 7E).

To gain a global view of shared regulatory modules across perturbation clusters, we organized DEGs into gene programs and determined the average effect size of each GP for each perturbation versus cytomix-only controls (**Fig. 3D**, Extended Data Fig. 6C, Table S9). GPs with the highest effect size continued to be centered around the ‘TGFβ-superfamily’ and ‘Inflammatory’ perturbation clusters, confirming the transcriptional impact of these perturbations.

Not only did we observe unexpected similarities in GP activity between dissimilar ligand families, but also divergent GP activation between similar ones. For instance, BMP2/4/6 and TGFβ1/2 downregulated GPs 1 and 5, while BTC, HBEGF, and EGF upregulated these GPs, indicating a shared ability of both ligand families to modulate these programs. Conversely, GP6 was upregulated by BMP2/6 but downregulated by BMP4.

### Niche factor perturbations elicit distinct and cell type-specific transcriptional programs

We next leveraged the single cell resolution of our atlas to investigate cell type-specific response across secreted ligands. Some perturbations led to the emergence of new cell populations, such as the M cell-like population in RANKL-treated organoids (**Fig. 4A**)^52^. Other perturbations elicited drastic changes in cell type composition. BMP2, 4, and 6 led to a decrease in the proportion of stem cells and increase in colonocytes, consistent with BMP’s role in restricting stem cell hyperproliferation ^13,53^ (**Fig. 4B**, Extended Data Fig. 8). On the other hand, inflammatory factors like IFNγ and TNFα led to a decrease in colonocyte proportion (**Fig 4B,C**), in line with observations in human diseased vs. healthy IBD patient samples^4^. EGF, HBEGF, IFNα, RANKL and overall cytomix stimulation decreased goblet cell proportions while IL22, BMP6, and TRAIL *(TNFSF10*) increased it. Not only has recent evidence suggested that increased goblet cell abundance is associated with clinical remission following treatment, goblet cells in TRAIL-tread organoids displayed increased expression of remission-associated goblet markers (*MUC2, SPINK4, LCN2*) (Table S8)^6^.

To further understand how DEGs are co-regulated in a cell-type specific manner, we calculated effect sizes of previously determined GPs (**Fig 3D**) across individual cell types for select high impact perturbations (**Fig. 4D**, Table S12). Cytomix treatment itself exerted the strongest effects, as no-cytomix organoids downregulated GP20 (immune response to cytokines: *IFITM3, IFI6, DDX60*) and GP13 (Neutrophil chemotaxis, response to cytokine: *CXCL9, CXCL10, CXCL11*), and upregulated GP2 (ribosome biogenesis: *RPS12, RPS28, RPL39*) across all cell types. GP7, related to organ growth and cell migration (*EDN1, ANXA3, WWC2*) was downregulated most strongly in differentiated cells with BMP2, 6, and 4 but not other perturbations. Interestingly, besides GP7, additional responses to BMP4 appeared to be more transcriptionally distinct from BMP2/6, suggesting divergences in the BMP family ligands (Extended Data Fig. 7A-D). GP6, related to proliferation and regulation of wound healing (*PTPRK, SH3RF2, HEPH*) exhibited opposing effects in BMP2/6 compared to EGF-family ligands, particularly in stem/TA cells. Notably, treatment with members of the TGFβ-superfamily, particularly BMPs, resulted in similar trends to no-cytomix organoids, suggesting that specific aspects of the non-inflamed epithelium can be recapitulated by administration of TGFβ-superfamily ligands in the presence of cytomix. Specifically, GP1 (Ion transport and cell adhesion: *PLCB4, ITPR2, SKAP1*), is downregulated in stem and TA cells in both no-cytomix and BMP2/6/TGFβ ligands and upregulated in TA cells for pro-inflammatory perturbations TNFα and IFNγ. In contrast, GP12 (epithelial structure maintenance and downregulation of anoikis: *FABP1, LGALS3, S100A14*) is upregulated in stem and/or TA cells in both no-cytomix and BMP2/6/TGFβ1 and downregulated in IL13, IL4, and pro-inflammatory perturbations **(Fig. 4D,E)**.

### Perturbation-elicited organoid gene programs are active in human IBD tissue

To probe the functional relevance of GPs elicited in organoids, we searched for their expression in human IBD tissue ^4,25^(**Fig. 5A**). We focused on GP1, 13, and 20 as they were upregulated by cytomix treatment and TNFα/ IFNγ individually, suggesting a pathogenic nature. These GPs additionally display large effect sizes in organoids upon stimulation with perturbations in the TGFβ-superfamily, IFNα, and IL4/13 (GP1 - downregulated by TGFβ-superfamily, GP13 - downregulated by IL4 and IFNα, GP20 - downregulated by BMP6 and IL13). Notably, many genes in these GPs showed a concordant decrease in healthy vs. IBD epithelium across two scRNA-seq datasets (**Fig. 5A**, Table S11). This suggests that increased expression of these programs may be associated with inflammation *in vivo* and that members of the TGFβ-superfamily, IFNα, and IL4/13 have the specific ability to modulate their expression. To examine the localization of these GPs in human tissue, we performed spatial transcriptomics (CosMx) on healthy (n=4) vs UC (n=16) patient samples (Table S10) (**Fig. 5B**). GP1 was excluded from this analysis due to low panel coverage. Not only were GP13 and GP20 epithelial-enriched, they were concordantly upregulated in UC vs. normal epithelium (**Fig. 5B,C**). Relative to all other cell types, GP20 was mostly increased in epithelial cells while GP13 was significantly upregulated across all compartments, suggesting that it is a pan-inflammatory signature (**Fig. 5C**). Lastly, we mapped the expression of GP1/GP13 and a previously-described fibroblast signature predictive of non-response to anti-TNFα therapy^54,55^ onto their respective cells types in human IBD scRNA-seq data^4^ and found them to be positively correlated at a patient level (Extended Data Fig. 9A). Altogether, this suggests that organoid-derived GPs are capable of modeling physiologically-relevant responses. In particular, we find that several GPs may be associated with a pathogenic inflammatory response, and members of the TGFβ-superfamily, IL4, IL13, and IFNα may play protective roles in the inflamed epithelium^56–60^.

Finally, we checked the expression of IBD-related GPs IBD1 and IBD2 **(Fig. 1**) across perturbations (**Fig. 5D,E**, Extended Data Fig. 9B, Table S12). Inflammation-associated IBD1 was slightly downregulated in TA cells from BMP6, IL13, and IL4-treated organoids^58^. We observed a stronger response for metabolism-associated IBD2, which was upregulated across the majority of cell types in no-cytomix along with BMP2/6, and in stem cells for TGFβ1/2/3 (**Fig. 5D,E)**. This further suggests that some members of the TGFβ-superfamily family of ligands as well as IL4/13 may recapitulate aspects of transcriptional states found in the uninflamed epithelium.

## DISCUSSION

Systematically charting microenvironmental regulation of the diseased intestinal epithelium is important for generating new therapeutic hypotheses and biomarkers in multifactorial diseases like IBD. We developed and characterized an organoid model reflecting IBD-relevant inflammatory properties and found that model selection is critical not only to recapitulate disease phenotypes, but impacts responses to niche cues in a dramatic manner. We then used this system to generate a scRNA-seq atlas of epithelial responses to diverse niche cues, where we uncovered previously unknown relationships between paracrine-mediated pathways in the gut epithelium. For instance, while many perturbations from shared ligand families showed similar patterns in expression of gene programs, some were divergent. Notably, GP6 was upregulated in BMP2, 5, and 6-treated organoids but downregulated in BMP4-treated organoids, highlighting the utility of comparing within ligand families.

Differences in activity could be due to differences in receptor affinity, as BMP6 binds preferentially to BMPR2 while BMP2/4 bind preferentially to BMPR1A/B^61^, or differences in induction of apolipoproteins^13^. Notably, many responses to niche cues are cell type specific. In particular, transit amplifying cells displayed large effect sizes for most GPs compared to other cell types, suggesting that these proliferative progenitors are more responsive to secreted cues, particularly in an inflamed context.

Of the perturbations that elicited a strong molecular response, members of the TGFβ-superfamily, particularly BMP2 and BMP6, were relatively more transcriptionally correlated with the ‘healthy’ control organoids that did not receive cytomix, indicating that these perturbations may play a role in epithelial homeostasis in the face of injury ^60^. However, only a subset of the DEGs elicited by BMP treatment overlapped with those found in control vs. cytomix organoids, indicating that BMPs only restore specific pathways out of many dysregulated by epithelial inflammation. Some of this similarity may be driven by metabolic shifts, as BMP2 and 6 show strong upregulation of metabolism-related GP IBD2.

Interestingly, many of the GPs we identified from the screen, particularly those displaying strong effects in no-cytomix organoids and upon treatment with TGFβs, BMPs, IL4, IL13, and IFNα, showed disease-relevance as they were also downregulated in healthy patients in both published scRNA-seq atlases and our spatial transcriptomic data. While these GPs show similar direction with healthy vs inflamed tissue, it is difficult to deconvolve which genes associated with inflamed tissue functionally modulate inflammation/stress vs. regeneration/repair. Such questions could be investigated by further refining the spatial analyses on human IBD tissue to distinguish GPs upregulated in sites of lesions vs. active repair.

Surprisingly, many perturbations did not yield a significant transcriptional response upon treatment of inflamed organoids. It is possible that such ligands exert context-specific responses (e.g. in development but not in disease), may require interactions with cell types other than the epithelium to elicit epithelial

## METHODS

### Generation of primary human colon organoids

Sections of human colon were provided by Donor Network West from deceased patients with no history of intestinal disease. To clean the tissue and isolate fresh crypts for organoid generation, colon tissue was first sliced open lengthwise and wiped 3-4 times with gauze (Fisher Scientific) to remove waste and mucus. The tissue was then cut into small sections of approximately 2” x 2” and placed into a 15cm dish filled with PBS++ (PBS supplemented with 10mM HEPES (Thermo Fisher Scientific), 1X GlutaMAX (Thermo Fisher Scientific), 1X Pen-Strep (Thermo Fisher Scientific), and 2.5 ug/mL Amphotericin B (Thermo Fisher Scientific)). The epithelium was then minced from tissue segments using scalpels (Fisher Scientific) and collected into 50mL conical tubes. Minced epithelium was washed 3 times with DMEM++ (Advanced DMEM/F12 (Thermo Fisher Scientific) supplemented with 10mM HEPES, 1X GlutaMAX, 1X Pen-Strep, and 2.5 ug/mL Amphotericin B), and centrifuged at 4C, 500g for 3 minutes between each wash. After the final wash, tissue was resuspended in 25mL of 2.5mM EDTA in PBS and placed in an end-over-end rotator at 37C until crypts were visible, around 12 minutes. Following digestion, the EDTA response which would not be evident in an epithelial-only organoid system, or require higher dosing than that which was utilized in the experiment.

We focused on epithelial-only organoids to cleanly assess the impact of secreted factors from supporting cell types on the epithelium. Future work could benefit from employing co-cultures of epithelium with stromal cells and/or immune cells to capture more complex synergistic interactions and feedback loops. While our work focused on recapitulating IBD-like phenotypes from healthy organoids, organoids derived directly from IBD patient tissue may bear even more physiological relevance^11^. However, because these disease phenotypes cannot be maintained *ex vivo* for an extended period of time^12^, identifying additional exogenous cues is necessary. Finally, the sample size in our study poses limitations, as only 3 donors were utilized. These findings would benefit from more donors to better capture patient diversity, particularly sex-specific differences. digestion solution was topped up with 20mL of DMEM++, filtered through sterile gauze, and centrifuged for 3 minutes at 4C, 500g. The pellet was then resuspended in 10mL PBS++, filtered through a 100μm cell strainer, and centrifuged for 3 minutes at 4C, 500. The tissue was then ready for plating in Matrigel (Corning) by resuspending in cold Matrigel and plating into 30uL Matrigel droplets, at a density of 30-50 crypts per droplet. Once Matrigel had solidified for 30 minutes at 37C, crypts were fed with organoid growth media^62^: Advanced DMEM/F12 supplemented with 1X Pen-Strep, 10mM HEPES, 1X GlutaMAX, 1X B-27 Supplement (Thermo Fisher Scientific), 0.2nM Wnt Surrogate protein (ImmunoPrecise Antibodies), 5nM Gastrin-1 (Tocris), 1.25mM N-Acetyl-L-Cysteine (Sigma-Aldrich), 10mM Nicotinamide (Sigma-Aldrich), 100μg/mL Primocin (InvivoGen), 500nM A 83-01 (Tocris), 50ng/mL recombinant human EGF (R&D Systems), 100ng/mL recombinant human Noggin (R&D Systems), 250ng/mL recombinant human R-spondin 3 (Roche), 100ng/mL recombinant human IGF1 (BioLegend), 50ng/mL recombinant human FGF2 (PeproTech).

### Organoid maintenance and passaging

After the initial organoid plating, cultures were maintained in Cultrex Basement Membrane ECM (70% Cultrex - PBS) (R&D Systems) domes. Organoids were cultured every 5-7 days as needed based on size and morphology. To passage, organoids were mechanically dislodged from Cultrex domes using a P1000 pipette tip and resuspended in cold 10mM EDTA with 10μM Y-27632 (STEMCELL Technologies) in a 15mL conical tube. The tube containing organoids was then placed on an end-over-end rotator at 4C for 30 minutes and then centrifuged for 5 minutes at 500g, 4C. Organoids were then resuspended in 1X Trypsin (Fisher Scientific) and placed in an end-over-end rotator for 6 minutes at 37C. Following dissociation, Trypsin was neutralized with DMEM containing 10% FBS, and organoids were centrifuged for 5 minutes at 500g, 4C. Media was aspirated and organoids were washed once with PBS, followed by centrifugation for 5 minutes at 300g, 4C. PBS was aspirated and organoids were resuspended in 70% Cultrex with PBS, and plated in 30uL droplets in 6 well plates. The plates were placed upside down in the incubator (37C, 5% CO2) for 30 minutes, after which organoid growth media with 10μM Y-27632 was added. Media was changed every other day to organoid growth media.

### Organoid culture in microwell plates

Experiments involving microwell plates utilized 500μm Gri3D plates (SUN bioscience). Prior to organoid seeding, storage buffer was aspirated from each well and 150uL of Advanced DMEM/F12 was added through the pipetting port. The plate was placed in the tissue culture incubator for at least 30 minutes.

Organoids that had been maintained in Cultrex domes for at least 5 days after passaging were removed from the ECM following the passaging protocol as above, but dissociated for 12 minutes in Trypsin rather than 6, or until organoids were largely single-cells. Trypsin was neutralized and organoids were washed as in the passaging protocol. Following the PBS wash, organoids were resuspended in organoid growth media without Nicotinamide: Advanced DMEM/F12 supplemented with 1X Pen-Strep, 10mM HEPES, 1X GlutaMAX, 1X B-27 Supplement, 0.2nM Wnt Surrogate protein, 5nM Gastrin-1, 1.25mM N-Acetyl-L-Cysteine, 100μg/mL Primocin, 500nM A 83-01, 50ng/mL recombinant human EGF, 100ng/mL recombinant human Noggin, 250ng/mL recombinant human R-spondin 3, 100ng/mL recombinant human IGF1, 50ng/mL recombinant human FGF2, and with 10μM Y-27632. Dissociated organoids were then counted using a hemocytometer and seeded into the microwell plates at a density of 200 cells per microwell, in 50μL of media per well. The plates were placed in the incubator for 30-60 minutes to allow the cells to settle, after which another 100μL of media containing 2% Matrigel was added per well. It is critical that Matrigel is added to the media while cold. On the day after seeding, media was removed and changed to growth media without Y-27632. On day 3 after seeding, media was removed and organoids were washed twice with Advanced DMEM/F12. After washing, media was replaced with organoid media without EGF to promote differentiation. Media was changed every 1-2 days, and cultures were maintained in the plates for up to 10 days before collection.

### Injury models in organoids

For experiments involving irradiation, DSS, and cytomix, organoids were cultured as above in microwell plates for 5 days. On the 5th day after seeding, injury conditions were applied. For irradiation studies, the plates were placed on the center of the rotating table of the irradiator (J.L. Shepherd and Associates, Mark I series, model 68 A, Cesium (137Cs) gamma irradiator) for the time periods equivalent to either 6 Gy or 12 Gy, and then the organoids were further cultured in the incubator. For DSS treatment, DSS (MP Biomedicals) was dissolved in organoid media (without EGF or Nicotinamide), and added to organoid wells overnight for 16 hours, after which organoids were washed twice with Advanced DMEM/F12 and then replaced with organoid media. The low and high doses considered were 0.1% DSS and 0.5% (wt/vol) DSS in media. For cytomix treatment, human recombinant TNFα (R&D Systems), IFNγ (R&D systems), and IL-1β (PeproTech) were dissolved in organoid media (without EGF or Nicotinamide) for 8 hours, after which organoids were washed twice with Advanced DMEM/F12 and then replaced with organoid media. The low and high doses considered were 10ng/mL TNFα, 10 ng/mL IFNγ, 1 ng/mL IL1β and 50ng/mL TNFα, 50 ng/mL IFNγ, 5 ng/mL IL1β. Experiments were conducted on the following day after injury (scRNA-seq, imaging) to assess response to injury.

### FITC-dextran and apoptosis measurements in organoids

Organoid permeability and apoptosis measurements were conducted on one day post-injury. FITC-dextran was used to assess barrier integrity similar to previously published methods ^34^. Briefly, organoids were incubated in a solution of 4kDA FITC-Dextran (MedChemExpress) diluted to 2mg/mL in PBS for 5 minutes in μ-slides (Ibidi) to allow organoids to settle to the bottom of the slide chamber. Organoids were then imaged on a Nikon spinning disc confocal to collect fluorescence and DIC images, and the percentage of intact organoids was calculated as the number of organoids per condition/donor that excluded FITC-dextran from the lumen divided by the total number of organoids. For apoptosis quantification, organoids were incubated in CellEvent Caspase-3/7 Detection Reagent (ThermoFisher) diluted in media according to manufacturer protocols for 30 minutes in the microwell plates. Organoids were then washed with PBS and imaged on a Nikon spinning disc confocal to collect fluorescence and DIC images. The % area of activated Caspase-3/7 was quantified using FIJI^63^ as the area of organoid stained positive for Caspase-3/7 divided by the total area of the organoid from DIC images.

### Organoid whole-mount immunofluorescence

For immunofluorescent images, organoids were removed from the microwell plates and fixed in 4% paraformaldehyde (ThermoFisher) overnight at 4C in 1.5mL Eppendorf tubes on a rocker. On the next day, organoids were washed three times with PBS and resuspended in permeabilization/block buffer (5% normal donkey serum (Abcam), 0.5% Triton X-100 (ThermoFisher) in PBS)^64^ overnight at 4C on a rocker. Organoids were then resuspended in primary antibody diluted in organoid wash buffer (0.1% Triton X-100, 0.2% BSA (Miltenyi)) overnight at 4C on a rocker. On the following day, organoids were washed three times (2 hours per wash at 4C on a rocker) in organoid wash buffer, and then resuspended in secondary antibody with DAPI diluted in organoid wash buffer. On the next day, organoids were washed three times with PBS (2 hours per wash at 4C on a rocker), and then placed in a 96-well glass bottom plate. PBS was carefully removed and organoids were resuspended in 60 uL of CUBIC-R (TCI). After allowing organoids to settle, imaging was conducted on a Leica STED. Brightness and contrast adjustments were performed using Fiji^63^, and adjustments were made uniformly across images.

### Single cell preparation of organoids for single cell RNA-sequencing

For scRNA-seq experiments, organoids were dissociated using the Neural Dissociation-P kit (Miltenyi). Organoids were pipetted out of their microwell plates and into a deep-well 96 well plate (ThermoFisher) for dissociation. Enzyme mix 1 was prepared according to manufacturer protocols with the addition of Y-27632 for a final volume of 1.5mL per well. Organoids in mix 1 were then placed at 10C for 10 minutes, after which enzyme mix 2 was added.

Organoids were kept at 10C throughout dissociation for up to 60 minutes, and were removed every 10 minutes to be pipette mixed with a P1000 pipette to aid in dissociation. Once dissociation was complete, organoids were centrifuged for 3 minutes, 500g, 4C and washed twice with PBS. For pooled experiments involving hashing (perturbation experiment, pooled injury experiments), organoids were resuspended in 100μL cell staining buffer (2% BSA, 0.01% Tween in PBS) and then blocked with 5uL human TruStain FCX blocking buffer (BioLegend) for 10 minutes on ice. 1μL of a unique cell hashing antibody (TotalSeqA: BioLegend) was added per condition for 20 minutes on ice, and cells were pipette mixed after 10 minutes.

Organoids were then washed three times with cell staining buffer on ice. After the final wash, organoids were resuspended in 0.04% BSA in PBS, filtered through a 70μm Flowmi cell strainer (Sigma Aldrich) and loaded onto a 10X Chromium X according to manufacturer instructions. For organoid and tissue characterization and pooled injury experiments, the standard 10x 3’ v3.1 kit was used. For some injury experiments, the 10x 3’ low throughput kit was used to compare multiple injuries from the same experiment. For the perturbation dataset, the 10x 3’ high throughput kit was used. Gene expression libraries were prepared according to 10X protocols, and HTO libraries were prepared according to published protocols^65^.

### scRNA-seq data pre-processing

Libraries were sequenced on a NextSeq2000 or NovaSeq6000 (for HT libraries) targeting 30,000 reads/cell for GEX libraries and 2,000 reads/cell for HTO libraries. Sequenced reads were aligned using the human reference genome GRCh38-2020-A using the CellRanger pipeline (10x Genomics). scRNA-seq data was processed using CellRanger and resulting count matrices were analyzed using Seurat in R. For experiments involving hashing, cells were demultiplexed using the HTODemux function in R. For experiments involving genetic demultiplexing as in the perturbation dataset, the souporcell algorithm^66^ was used to distinguish between organoids from unique donors. Doublets were removed from data using the DoubletFinder algorithm ^67^. When combining experiments from multiple donors or timepoints, data was batch-corrected using the Harmony algorithm^68^.

### Secreted niche factor perturbation screen

Human colon organoids from 3 independent donors were expanded in Cultrex domes and then seeded in microwell plates as previously described in a matched experiment. On the 5th day post-seeding, low-dose cytomix (10ng/mL TNFα, 10 ng/mL IFNγ, 1 ng/mL IL1β) was applied to the organoids in a media change with 0.5% Matrigel. At the time of cytomix addition, human recombinant proteins corresponding to the 81 selected perturbations (Table S14) were applied to each well in an arrayed format. After 8 hours, cytomix was removed from the organoids by washing twice with Advanced DMEM/F12 for 30 minutes in the incubator. After washing, fresh organoid media with 0.5% Matrigel and human recombinant proteins was added to each well (Table S14). Media was changed every 1-2 days with fresh organoid media containing human recombinant proteins, and organoids were collected for scRNA-seq on day 8 post-seeding. Dissociated organoids were loaded onto 4 3’ HT channels (10X), and for each independent channel one no-cytomix control as well as 3 cytomix-only controls were included.

### Spatial Transcriptomics (CosMx) data collection for human colon tissue

Formalin-fixed, paraffin-embedded (FFPE) tissue blocks were sectioned at 5 μm thickness using a microtome and processed according to Nanostring’s protocol for “CosMx SMI Semi-Automated Slide Preparation for RNA Assays”. Briefly, the slides were placed in an oven at 60°C overnight to facilitate paraffin removal and sample conditioning.

Deparaffinization and antigen retrieval were performed using the Leica Bond RX system. Slides were incubated with ER1 (AR9961, Leica) at 100°C for 20 minutes to retrieve antigens. Subsequently, proteinase K (AM2546, ThermoFisher) was applied at a concentration of 3 μg/mL for 15 minutes to enhance tissue permeability and facilitate probe hybridization. Fiducial markers were applied for 5 minutes to aid in downstream imaging. Afterwards, slides were incubated with 100 mM NHS-acetate for 15 minutes to block nonspecific binding sites, followed by washing with 2X SSC buffer. Post-fixation was carried out using 10% neutral buffered formalin (16004-122, VWR) for 5 minutes. Subsequently, slides were incubated with 100 mM NHS-acetate for 15 minutes to block nonspecific binding sites, followed by washing with 2X SSC buffer. Denatured CosMx RNA probe mixes were applied to the slides. Hybridization was performed overnight under nuclease-free conditions (RNase away and DEPC treated water) in a hybridization oven (HybEZ, ACD) at 37°C.

After hybridization, stringent washes were performed twice using a stringent solution (50% formamide in 2X SSC buffer solution) at 37°C for 25 minutes each.

Unbound probes were removed and background signal reduced by washes with 2X SSC buffer solution. Slides were incubated with nuclear stain for 15 minutes, followed by application of cell segmentation mix (CD298/B2M) with supplemental markers (PanCK/CD45) for 1 hour. Slides were washed with 1X PBS to remove excess staining reagents and flow cells were assembled to the slides. To prevent tissue desiccation, 2X SSC buffer was pipetted into the flow cell ports.

The prepared slides and the CosMx RNA imaging 1000-plex tray were loaded onto the CosMx Spatial Molecular Imaging (SMI) platform. The flow cell setting was created for pre-bleaching profile using Colorectal Tissue (configuration A) and the cell segmentation profile used was Human (configuration A). Scan area and fields of view (FOV) were selected for each slide and data acquired during the cycling phase on the CosMx (SMI) platform, capturing high-resolution, spatially resolved RNA expression profiles. During the CosMx cycling, data was automatically transferred to the AtoMx platform for further analysis.

## COMPUTATIONAL METHODS

### Cell type annotation in organoids

Organoid scRNA-seq data was quality controlled, filtered and annotated using Seurat (v5.0.1). Cell-type annotation was performed by applying FindTransferAnchors and TransferData using our matched colonic tissue reference atlas (Extended Data Fig. 1), and then manually cleaning up annotations based on expression of canonical cell type markers. The human colon organoid tissue atlas was first annotated by integrating our colon data with published atlases containing healthy colon tissue ^4,25,69^ using scANVI to harmonize cell type labels in a consistent manner, thus generating an integrated atlas.

### Derivation of colon epithelial IBD signatures from published atlases

scRNA-seq data downloaded from public datasets (Smillie et al., Kong et al., and Parikh et al.)^4,25,46^ were imported and cells with greater than 30% mitochondrial gene content and fewer than 200 genes expressed were removed from all datasets for downstream analysis. Cell-type annotation was re-performed by projecting each query dataset onto our human colon integrated atlas (using Seurat functions FindTransferAnchors and TransferData). Filtered RNA counts matrices were converted to h5ad format via the scanpy (v1.9.3) for input into consensus non-negative matrix factorization (cNMF). The cNMF package (v1.3.4) was utilized to identify gene modules with a number of components (k) set to 10, a local density threshold of 0.2, and a maximum of 10 iterations.

Meta-modules were identified by calculating the Spearman’s correlation using the gene spectra score space of cNMF modules derived independently from Smillie and Kong datasets. The 50 genes with greatest spectra scores from the modules within each meta-module were identified and the unions of these gene sets were the IBD1 and IBD2 colon epithelial IBD signatures.

For both the effect-size dotplots and the boxplots visualizing enrichment of metamodules in patients with inflamed vs non-inflamed conditions, IBD1/IBD2 signature scores were computed using the AddModuleScore function in Seurat. The means of module scores for each patient were then calculated. Effect sizes were evaluated using Cohen’s d formula for inflamed and non-inflamed groups for each cell type. Statistical significance between these groups was assessed using a t-test.

The predictive power of IBD1 and IBD2 for disease status was assessed by training a generalized linear model (GLM) on the Kong et al. dataset^25^, with 70% of the data used for training and testing on the remaining 30%, and finally validating in Smillie et al. dataset^4^.

### Cosine similarity, DEG, gene program and effect size analyses

Cosine similarities between perturbations was calculated on the Harmony donor-corrected PCA space (first 50 PCs). An average PC coordinate was calculated for each perturbation. To calculate the ‘donor-donor correlation’, average PC coordinates were calculated on a donor basis and cosine similarities were calculated for each donor-perturbation pairwise comparison. Significance was met if the average donor-donor correlation for a single perturbation was > 95% percentile of a permuted distribution (rho=0.406).

For the injury dataset, DEGs were calculated for each injury vs. control for each cell type, by fitting the data to a gamma-possion model via the glmGamPoi package using only the top 5000 highly variable genes and a randomly downsampled 300 cells per cell type per injury. LogFC values were shrunken. For the perturbation dataset, DEGs were calculated by pseudobulking counts for each gene on a donor basis and by cell types. DEseq2 was used to compare each perturbation to the untreated control, with the design testing for ‘perturbation’ while controlling for ‘donor.’ DEGs were designated as padj<0.05 and log2FC>0.2. ‘Average number of DEGs’ for each perturbation was calculated as the average across each major cell type for each perturbation.

To group perturbation-elicited DEGs into biologically interpretable gene programs, we ran cNMF^70^ on all DEGs in each dataset with a number of components (k) et to 20, a local density threshold of 0.2, and a maximum of 10 iterations. The resulting module usages were utilized for analyses comparing GP expression. ‘Activity’ vs. ‘identity’ modules were manually selected based on expression of canonical cell type-specific markers and restriction of module expression to particular cell types, and only cell activity modules were considered for further evaluation.

In the 81-perturbation data set, the effect size was calculated using the module usage scores (from cNMF), and each perturbation (3 donors, 1 well per donor) was only compared against the controls (3 donors, 3 wells per donor) that were from the same cDNA/library pool, to control for pool effects. The standard deviation and mean was taken across 3 donors for each perturbation. The p-value for each effect size was calculated using a t-test on the average module usage scores on a well/donor-basis, and FDR corrected.

Effect sizes of IBD1 and IBD2 were calculated by approximating the average expression of genes in each GP with AddModuleScore in Seurat, considering the top 50 genes in each GP based on gene score. For the injury dataset, the mean/standard deviation was taken across metacells computed across the dataset using the hdWGCNA package^71^ in R, with k=20 and max_shared = 4.

### Gene Ontology analysis: pathway enrichment of selected gene programs

GO analysis was performed using the enrichR package ^72^ in R, specifically querying the GO biological processes database from 2023. The top 50 genes from each GP based on gene spectra score were selected for inclusion in GO analysis.

### Selection of secreted factors for perturbation screen through ligand-receptor analysis

The perturbation space was established by mining through published scRNA-seq atlases of human colon from development (Yu et al., Elmentaite et al.)^26,27^, adult homeostasis (Elmentaite et al., Kong et al.)^25,26^, and IBD (Kong et al.) ^25^. Data from each atlas was considered individually and filtered to only include samples from the colon. Ligand-receptor interactions were assessed in each atlas using CellChat^73^, with trim = 0.02 and min.cells = 10. From each dataset, perturbations signaling to the epithelium were selected by setting targets.use to each epithelial cell type as annotated in the original dataset. After running CellChat on each dataset, perturbation lists were combined into a spreadsheet documenting each predicted signaling interaction and the probability of this interaction (Table S6), and prediction probabilities were averaged across all predicted interactions across datasets. The perturbation space was cut down to 81 to fit onto one 96-well plate (arrayed format) including controls by removing low-probability perturbations, perturbations exclusively signaling from immune cells, and preferentially selecting or adding in ligands from the same family or with known biological function (IFNα, IL1α, BMP3, DKK1, EdIL3, IFNE, IL4, SLIT3, ISLR, IL22).

### Calculation of logFC and p values in external datasets

For external IBD datasets^4,25^, DEGs were calculated by pseudobulking counts for each gene on a donor basis and by cell types. DEseq2 was used to compare healthy vs. diseased cells, with the design testing for ‘disease state’ while controlling for ‘donor’.

### Cell type composition analysis

Changes in cell type composition upon perturbations compared to the ‘Control’ were determined using a beta-binomial regression framework via the DCATS package^74^, with n=3 donors per perturbation. Results were additionally confirmed via differential abundance testing via MILO^75^ by considering donor and pool as covariates.

### Spatial Transcriptomics (CosMx) analysis

Analysis of spatial transcriptomics data generated by the CosMx SMI 1000-plex protocol was performed in two python workflows following standard quality control procedures output by the AtoMx segmentation and transcript assignment: 1) ensemble variational inference and label transfer and 2) gene set analysis and visualization.

First, ensemble variational inference and label transfer was used to annotate segmented single-cell spatial transcriptomes at a high resolution using a publicly available scRNA-seq datasets as a reference^37^. We performed 10 replicate runs of scVI followed by scANVI as a Papermill workflow on an Nvidia Tesla V100 32GB GPU. scVI parameters were as follows: n_latent = 32; max_epochs_scvi = 100; dropout rate = 0.3 (for resection samples) and 0.1 (for TMA samples); and batch_key = “Modality”, where we encoded “Reference” or “Spatial query” corresponding to the appropriate datasets. scANVI parameters were as follows: n_samples_per_label: 1000. In order to ensemble the results of all replicate runs we applied a classification entropy filter followed by consensus voting. Each replicate has an associated cell ID by cell type classification probability matrix, where the cell type classification entropy was calculated cell-wise and thresholded by finding the lowest local minimum by examining the histogram. Cells with classification entropies above this threshold are considered “Unknowns”, interpreted as mixed single-cell transcriptomes that could not be confidently assigned to a single cell type. Next, consensus voting is performed and the final annotation is assigned by majority vote given a proportional threshold of .5, where classifications below this number are again assigned as unknowns. For cell type-oriented analyses these unknowns were excluded. Further lineage-specific cell type refinements were performed through high-resolution leiden clustering on the output scANVI embedding.

Second, spatial gene set analysis was performed using the scanpy and squidpy python libraries. These data were normalized by library size and log1p transformed before scoring using the scanpy score_genes function with default parameters on each single cell. Gene set scores were calculated only on the intersection of the 1000-plex CosMx panel. For GP analyses in the context of the TMA samples, this scoring procedure was performed and averaged per biological replicate and per cell type. Mann-Whitney U tests were performed to assess statistical significance. For GP analyses in the context of the resected normal and UC samples, the same scoring procedure was performed; to visualize, the resulting scores were binned into 20 quantiles and plotted by their assigned 2D spatial coordinates.

## Data Availability

Data related to this manuscript will be deposited upon publication.

## Code Availability

Code related to this manuscript will be deposited upon publication.

## Acknowledgements

We are grateful for the cooperation of Donor Network West and all of the organ and tissue donors and their families, for giving the gift of life and the gift of knowledge by their generous donation. Additionally, we are thankful for Leslie Gaffney at the Broad Research Communication Lab for advice and editing of figures, and for Neko Ota for his help with organoid irradiation.

## Author contributions

MMC and MBC conceived the study. MMC performed all experiments. MMC, MBC, BC, KA, and RW analyzed all experiments. EP, VIL, CF, SR, and LMG performed all spatial transcriptomics experiments. LH, MMC, DP, ES, MK, and MBC contributed to organoid generation from human colon tissue and generation of organoid and tissue scRNA-seq data. XT and JL conceived of organoid cytomix treatment. CS performed organoid microscopy. MMC and MBC wrote the manuscript and produced the figures. All of the authors edited the manuscript.

## REFERENCES

1. Clevers, H. The Intestinal Crypt, A Prototype Stem Cell Compartment. Cell 154, 274–284 (2013).

2. Abud, H. E., Amarasinghe, S. L., Micati, D. & Jardé, T. Stromal Niche Signals That Orchestrate Intestinal Regeneration. CMGH 17, 679–685 (2024).

3. Abraham, C. & Cho, J. H. Inflammatory Bowel Disease. N Engl J Med 361, 2066–2078 (2009).

4. Smillie, C. S. et al. Intra- and Inter-cellular Rewiring of the Human Colon during Ulcerative Colitis. Cell 178, 714-730.e22 (2019).

5. Kinchen, J. et al. Structural Remodeling of the Human Colonic Mesenchyme in Inflammatory Bowel Disease. Cell 175, 372-386.e17 (2018).

6. Thomas, T. et al. A longitudinal single-cell atlas of anti-tumour necrosis factor treatment in inflammatory bowel disease. Nature Immunology 25, pages2152– 2165 (2024).

7. Holloway, E. M. et al. Mapping Development of the Human Intestinal Niche at Single-Cell Resolution. Cell Stem Cell 28, 568-580.e4 (2021).

8. Wen, C., Chen, D., Zhong, R. & Peng, X. Animal models of inflammatory bowel disease: category and evaluation indexes. Gastroenterology Report 12, goae021 (2024).

9. Kiesler, P., Fuss, I. J. & Strober, W. Experimental Models of Inflammatory Bowel Diseases. CMGH 1, 154–170 (2015).

10. Ferreira, B. et al. Trends in 3D models of inflammatory bowel disease. Biochimica et Biophysica Acta (BBA) - Molecular Basis of Disease 1870, 167042 (2024).

11. Meir, M. et al. Enteroids Generated from Patients with Severe Inflammation in Crohn’s Disease Maintain Alterations of Junctional Proteins. J Crohns Colitis 14, 1473–1487 (2020).

12. Arnauts, K. et al. Ex Vivo Mimicking of Inflammation in Organoids Derived From Patients With Ulcerative Colitis. Gastroenterology 159, 1564–167 (2020).

13. Beumer, J. et al. BMP gradient along the intestinal villus axis controls zonated enterocyte and goblet cell states. Cell Reports 38, 110438 (2022).

14. Chen, L. et al. TGFB1 induces fetal reprogramming and enhances intestinal regeneration. Cell Stem Cell 30, 1520-1537.e8 (2023).

15. Lukonin, I. et al. Phenotypic landscape of intestinal organoid regeneration. Nature 586, 275–280 (2020).

16. Zhang, J. et al. Tahoe-100M: A Giga-Scale Single-Cell Perturbation Atlas for Context-Dependent Gene Function and Cellular Modeling. bioRxiv (2025).

17. Sanchís-Calleja, V. O. P. et al. Decoding morphogen patterning of human neural organoids with a multiplexed single-cell transcriptomic screen. bioRxiv (2024).

18. Cui, A. et al. Dictionary of immune responses to cytokines at single-cell resolution. Nature 625, 377– 384 (2023).

19. Ayyaz, A. et al. Single-cell transcriptomes of the regenerating intestine reveal a revival stem cell. Nature 121–125 (2019).

20. Orzechowska-Licari, E. J., LaComb, J. F., Giarrizzo, M., Yang, V. W. & Bialkowska, A. B. Intestinal Epithelial Regeneration in Response to Ionizing Irradiation. J Vis Exp 185, (2023).

21. Montenegro-Miranda, P. S. et al. A Novel Organoid Model of Damage and Repair Identifies HNF4α as a Critical Regulator of Intestinal Epithelial Regeneration. Cellular and Molecular Gastroenterology and Hepatology 10, 209–223 (2020).

22. Katsandegwaza, B., Horsnell, W. & Smith, K. Inflammatory Bowel Disease: A Review of Pre-Clinical Murine Models of Human Disease. Int J Mol Sci 23, 9344 (2022).

23. Macedo, M. H., Neto, M. D., Pastrana, L., Gonçalves, C. & Xavier, M. Recent Advances in Cell-Based In Vitro Models to Recreate Human Intestinal Inflammation. Advanced Science 10, 2301391 (2023).

24. Aherne, C. M. et al. Neuronal guidance molecule netrin-1 attenuates inflammatory cell trafficking during acute experimental colitis. Gut 61, 695–705 (2012).

25. Kong, L. et al. The landscape of immune dysregulation in Crohn’s disease revealed through single-cell transcriptomic profiling in the ileum and colon. Immunity 56, 444-458.e5 (2023).

26. Elmentaite, R. et al. Cells of the human intestinal tract mapped across space and time. Nature 597, 250– 255 (2021).

27. Yu, Q. et al. Charting human development using a multi-endodermal organ atlas and organoid models. Cell 184, 3281-3298.e22 (2021).

28. Brandenberg, N. et al. High-throughput automated organoid culture via stem-cell aggregation in microcavity arrays. Nature Biomedical Engineering 863–874 (2020).

29. Wang, Q. et al. Cytokine-induced epithelial permeability changes are regulated by the activation of the p38 mitogen-activated protein kinase pathway in cultured Caco-2 cells. Shock 29, 531–7 (2008).

30. Antoni, L., Nuding, S., Wehkamp, J. & Stange, E. F. Intestinal barrier in inflammatory bowel disease. World J Gastroenterol 20, 1165–1179 (2014).

31. Iwamoto, M., Koji, A., Makiyama, K., Kobayashi, N. & Nakane, P. K. Apoptosis of Crypt Epithelial Cells in Ulcerative Colitis. Journal of Pathology 180, 152–159 (1996).

32. Michielan, A. & D’Incà, R. Intestinal Permeability in Inflammatory Bowel Disease: Pathogenesis, Clinical Evaluation, and Therapy of Leaky Gut. Mediators Inflamm 628157 (2015).

33. d’Aldebert, E. et al. Characterization of Human Colon Organoids From Inflammatory Bowel Disease Patients. Front Cell Dev Biol 8, (2020).

34. Co, J. Y. et al. Controlling Epithelial Polarity: A Human Enteroid Model for Host-Pathogen Interactions. Cell Reports 26, 2509-2520.e4 (2019).

35. Oost, K. C. et al. Dynamics and plasticity of stem cells in the regenerating human colonic epithelium. bioRxiv (2023).

36. Gregorieff, A., Liu, Y., Inanlou, M. R., Khomchuk, Y. & Wrana, J. L. Yap-dependent reprogramming of Lgr5+ stem cells drives intestinal regeneration and cancer. Nature 526, 715–718 (2015).

37. Li, J. et al. Identification and multimodal characterization of a specialized epithelial cell type associated with Crohn’s disease. Nature Communications 15, (2024).

38. Grasberger, H. et al. DUOX2 variants associate with preclinical disturbances in microbiota-immune homeostasis and increased inflammatory bowel disease risk. J Clin Invest 131, e141676 (2021).

39. Bolton, C. et al. An Integrated Taxonomy for Monogenic Inflammatory Bowel Disease. Gastroenterology 162, 859–876 (2022).

40. Song, F. et al. The role of alcohol dehydrogenase 1C in regulating inflammatory responses in ulcerative colitis. Biochem Pharmacol 192, 114691 (2021).

41. Santhanam, S. et al. Mitochondrial electron transport chain complex dysfunction in the colonic mucosa in ulcerative colitis. Inflamm Bowel Dis 18, 2158–68 (2012).

42. MacFie, T. S. et al. DUOX2 and DUOXA2 form the predominant enzyme system capable of producing the reactive oxygen species H2O2 in active ulcerative colitis and are modulated by 5-aminosalicylic acid. Inflamm Bowel Dis 20, 514–24 (2014).

43. Vatn, S. S. et al. Mucosal Gene Transcript Signatures in Treatment Naïve Inflammatory Bowel Disease: A Comparative Analysis of Disease to Symptomatic and Healthy Controls in the European IBD-Character Cohort. Clin Exp Gastroenterol 5–25 (2022).

44. Schniers, A. et al. Ulcerative colitis: functional analysis of the in-depth proteome. Clin Proteomics 16, (2019).

45. Barnich, N. et al. CEACAM6 acts as a receptor for adherent-invasive E. coli, supporting ileal mucosa colonization in Crohn disease. J Clin Invest 117, 1566– 1574 (2007).

46. Parikh, K. et al. Colonic epithelial cell diversity in health and inflammatory bowel disease. Nature 567, 49–55 (2019).

47. Abud, H. E., Chan, W. H. & Jardé, T. Source and Impact of the EGF Family of Ligands on Intestinal Stem Cells. Front Cell Dev Biol 9, 685665 (2021).

48. Keir, M., Yi, T., Lu, T. & Ghilardi, N. The role of IL-22 in intestinal health and disease. J Exp Med 217, e20192195 (2020).

49. Wagner, F. et al. Dose escalation randomised study of efmarodocokin alfa in healthy volunteers and patients with ulcerative colitis. Gut 72, 1451–1461 (2023).

50. Koch, S. et al. The Wnt antagonist Dkk1 regulates intestinal epithelial homeostasis and wound repair. Gastroenterology 141, 259–68, 268.e1–8 (2011).

51. Xu, J. et al. Secreted stromal protein ISLR promotes intestinal regeneration by suppressing epithelial Hippo signaling. EMBO J 39, e103255 (2020).

52. Lau, W. de et al. Peyer’s Patch M Cells Derived from Lgr5+ Stem Cells Require SpiB and Are Induced by RankL in Cultured “Miniguts.” Mol Cell Biol 32, 3639–3647 (2012).

53. Qi, Z. et al. BMP restricts stemness of intestinal Lgr5+ stem cells by directly suppressing their signature genes. Nature Communications 8, 13824 (2017).

54. Friedrich, M. et al. IL-1-driven stromal–neutrophil interactions define a subset of patients with inflammatory bowel disease that does not respond to therapies. Nature Medicine 27, 1970–1981 (2021).

55. Martin, J. C. et al. Single-Cell Analysis of Crohn’s Disease Lesions Identifies a Pathogenic Cellular Module Associated with Resistance to Anti-TNF Therapy. Cell 178, 1493-1508.e20 (2019).

56. Wirtz, S. & Neurath, M. F. Illuminating the role of type I IFNs in colitis. JCI 115, 586–588 (2005).

57. Li, J.-Y. et al. IRF/Type I IFN signaling serves as a valuable therapeutic target in the pathogenesis of inflammatory bowel disease. International Immunopharmacology 92, 107350 (2021).

58. Giuffrida, P., Caprioli, F., Facciotti, F. & Sabatino, A. D. The role of interleukin-13 in chronic inflammatory intestinal disorders. Autoimmunity Reviews 18, 549– 555 (2019).

59. Miyoshi, H., Ajima, R., Luo, C. T., Yamaguchi, T. P. & Stappenbeck, T. S. Wnt5a Potentiates TGF-β Signaling to Promote Colonic Crypt Regeneration After Tissue Injury. Science 338, 108–113 (2012).

60. Xie, Z. et al. Recent developments on BMPs and their antagonists in inflammatory bowel diseases. Cell Death Discovery 9, (2023).

61. Sanchez-Duffhues, G., Williams, E., Goumans, M.-J.Heldin, C.-H. & Dijke, P. ten. Bone morphogenetic protein receptors: Structure, function and targeting by selective small molecule kinase inhibitors. Bone 138, 115472 (2020).

62. Fujii, M. et al. Human Intestinal Organoids Maintain Self-Renewal Capacity and Cellular Diversity in Niche-Inspired Culture Condition. Cell Stem Cell 23, 787-793.e6 (2018).

63. Schindelin, J. et al. Fiji: an open-source platform for biological-image analysis. Nature Methods 9, 676–682 (2012).

64. Childs, C. J. et al. EPIREGULIN creates a developmental niche for spatially organized human intestinal enteroids. JCI Insight 8, e165566 (2023).

65. Stoeckius, M. et al. Cell Hashing with barcoded antibodies enables multiplexing and doublet detection for single cell genomics. Genome Biology 19, (2018).

66. Heaton, H. et al. Souporcell: robust clustering of single-cell RNA-seq data by genotype without reference genotypes. Nature Methods 17, 615–620 (2020).

67. McGinnis, C. S., Murrow, L. M. & Gartner, Z. J. DoubletFinder: Doublet Detection in Single-Cell RNA Sequencing Data Using Artificial Nearest Neighbors. Cell Systems 8, 329-337.e4 (2019).

68. Korsunsky, I. et al. Fast, sensitive and accurate integration of single-cell data with Harmony. Nature Methods 16, 1289–1296 (2019).

69. Chen, B. et al. Differential pre-malignant programs and microenvironment chart distinct paths to malignancy in human colorectal polyps. Cell 184, 6262-6280.e26 (2021).

70. Kotliar, D. et al. Identifying gene expression programs of cell-type identity and cellular activity with single-cell RNA-Seq. eLife 8, e43803 (2019).

71. Morabito, S., Reese, F., Rahimzadeh, N., Miyoshi, E. & Swarup, V. hdWGCNA identifies co-expression networks in high-dimensional transcriptomics data. Cell Reports Methods 3, 100498 (2023).

72. Kuleshov, M. V. et al. Enrichr: A Comprehensive Gene Set Enrichment Analysis Web Server 2016 Update. Nucleic Acids Research 44, W90–W97 (2016).

73. Jin, S. et al. Inference and analysis of cell-cell communication using CellChat. Nature Communications 12, 1088 (2021).

74. Lin, X., Chau, C., Ma, K., Huang, Y. & Ho, J. W. K. DCATS: differential composition analysis for flexible single-cell experimental designs. Genome Biology 24, (2023).

75. Dann, E., Henderson, N. C., Teichmann, S. A., Morgan, M. D. & Marioni, J. C. Differential abundance testing on single-cell data using k-nearest neighbor graphs. Nature Biotechnology 40, 245–253 (2021).

76. Hao, Y. et al. Dictionary learning for integrative, multimodal and scalable single-cell analysis. Nature Biotechnology 42, 293–304 (2023).

77. Xu, C. et al. Probabilistic harmonization and annotation ofsingle-cell transcriptomics data with deepgenerative models. Molecular Systems Biology 17, e9620 (2021).

78. Love, M. I., Huber, W. & Anders, S. Moderated estimation of fold change and dispersion for RNA-seq data with DESeq2. Genome Biology 15, (2014).

