## Supplemental Figures for "Dictionary of human intestinal organoid responses to secreted niche factors at single cell resolution"

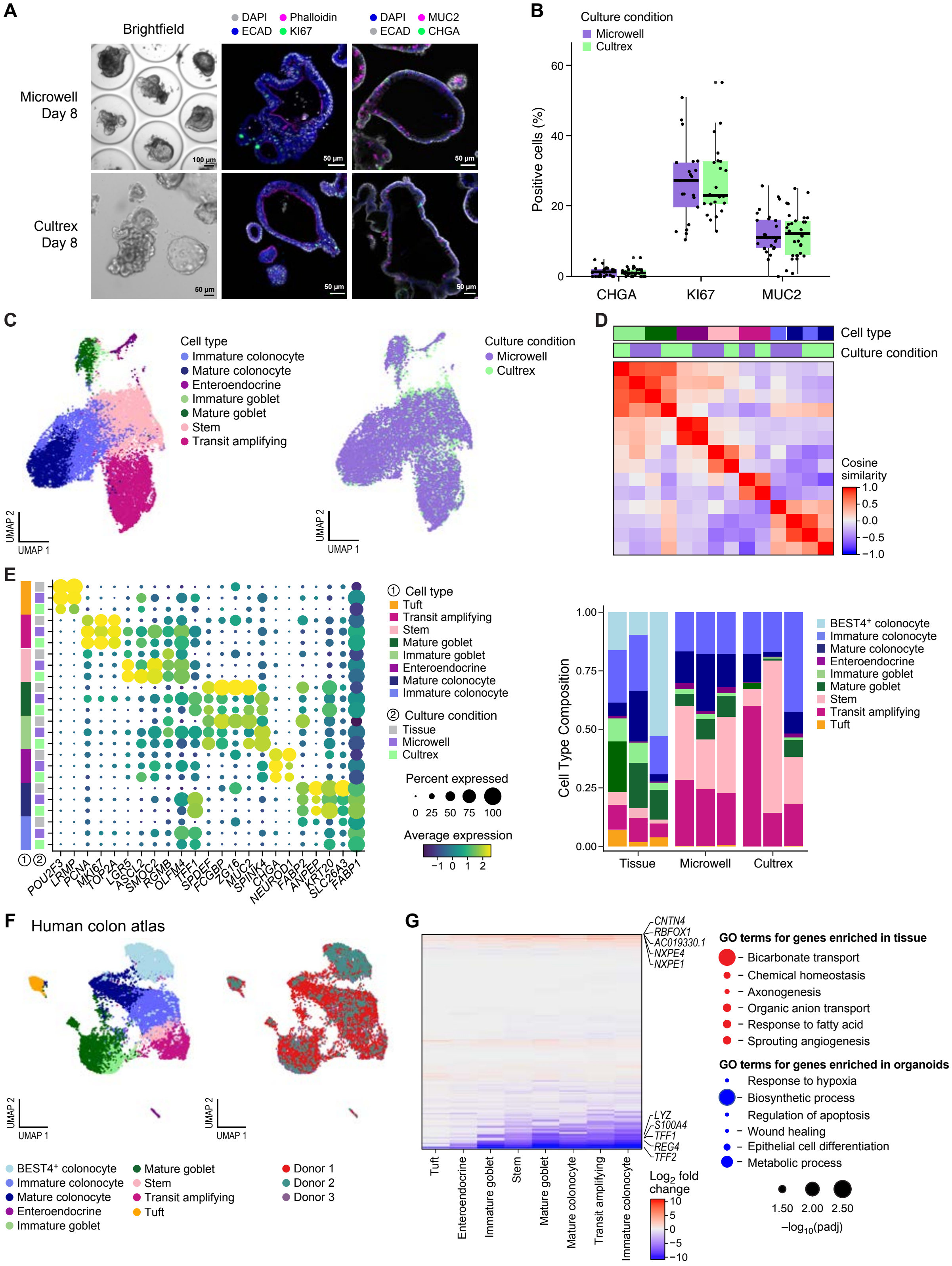
A
Brightfield
DAPI
Phalloidin
ECAD
KI67
DAPI
MUC2
ECAD
CHGA
Microwell
Day 8
Cultrex
Day 8
100 µm
50 µm
B
Culture condition
Microwell
Cultrex
Positive cells (%)
60
40
20
0
CHGA
KI67
MUC2
C
Cell type
Immature colonocyte
Mature colonocyte
Enteroendocrine
Immature goblet
Mature goblet
Stem
Transit amplifying
Culture condition
Microwell
Cultrex
UMAP 2
UMAP 1
D
Cell type
Culture condition
Cosine similarity
1.0
0.5
0.0
−0.5
−1.0
E
① Cell type
Tuft
Transit amplifying
Stem
Mature goblet
Immature goblet
Enteroendocrine
Mature colonocyte
Immature colonocyte
② Culture condition
Tissue
Microwell
Cultrex
Percent expressed
0 25 50 75 100
Average expression
−1 0 1 2
POU2F3
LRMP
PCNA
MKI67
TOP2A
LGR5
ASCL2
SMOC2
RGMB
OLFM4
TFF1
SPDEF
FCGBP
ZG16
MUC2
SPINK4
CHGA
NEUROD1
FABP2
ANPEP
KRT20
SLC26A3
FABP1
Cell Type Composition
1.00
0.75
0.50
0.25
0.00
Tissue
Microwell
Cultrex
BEST4+ colonocyte
Immature colonocyte
Mature colonocyte
Enteroendocrine
Immature goblet
Mature goblet
Stem
Transit amplifying
Tuft
F
Human colon atlas
UMAP 2
UMAP 1
BEST4+ colonocyte
Immature colonocyte
Mature colonocyte
Enteroendocrine
Immature goblet
Mature goblet
Stem
Transit amplifying
Tuft
Donor 1
Donor 2
Donor 3
G
CNTN4
RBFOX1
AC019330.1
NXPE4
NXPE1
LYZ
S100A4
TFF1
REG4
TFF2
Tuft
Enteroendocrine
Immature goblet
Stem
Mature goblet
Mature colonocyte
Transit amplifying
Immature colonocyte
Log2 fold change
10
5
0
−5
−10
GO terms for genes enriched in tissue
Bicarbonate transport
Chemical homeostasis
Axonogenesis
Organic anion transport
Response to fatty acid
Sprouting angiogenesis
GO terms for genes enriched in organoids
Response to hypoxia
Biosynthetic process
Regulation of apoptosis
Wound healing
Epithelial cell differentiation
Metabolic process
1.50 2.00 2.50
−log10(padj)

***Extended Data Figure 1: Human colon organoids cultured in microwell plates resemble gold-standard Cultrex-grown organoids and human colon tissue*** **(A)** Brightfield and immunofluorescent images (DAPI, Phalloidin, ECAD, KI67, MUC2, CHGA) comparing human colon organoids cultured for 8 days in standard Cultrex ECM domes to organoids cultured in microwell plates. **(B)** Quantification of organoid IF images (CHGA, KI67, and MUC2), n>= 20 organoids per condition. **(C)** Dimensional reduction of organoid scRNA-seq data from 3 donors/experiments of organoids cultured in Cultrex vs. microwell plates. **(D)** Cosine similarity between cell types from organoids cultured in Cultrex vs. microwell plates. **(E)** Left: Scaled expression of canonical cell type markers in Cultrex organoids, microwell organoids, and human colon tissue. Right: Celltype composition of Cultrex organoids, microwell organoids, and human colon tissue. Different bars represent different donors (n=3). **(F)** scRNA-seq data from three pathologically normal human colon donors. Plots depict cell type composition and donor heterogeneity. **(G)** Left: Log2 fold change of differentially expressed genes between tissue vs. organoid (red = upregulated in tissue, blue = upregulated in organoid) within each cell type. Right: GO terms associated with genes upregulated in tissue (red) or organoid (blue). DEGs were selected for inclusion in GO analysis based on padj <= 0.05 and abs(logFC) >= 0.5 from the union of DEGs across cell types.

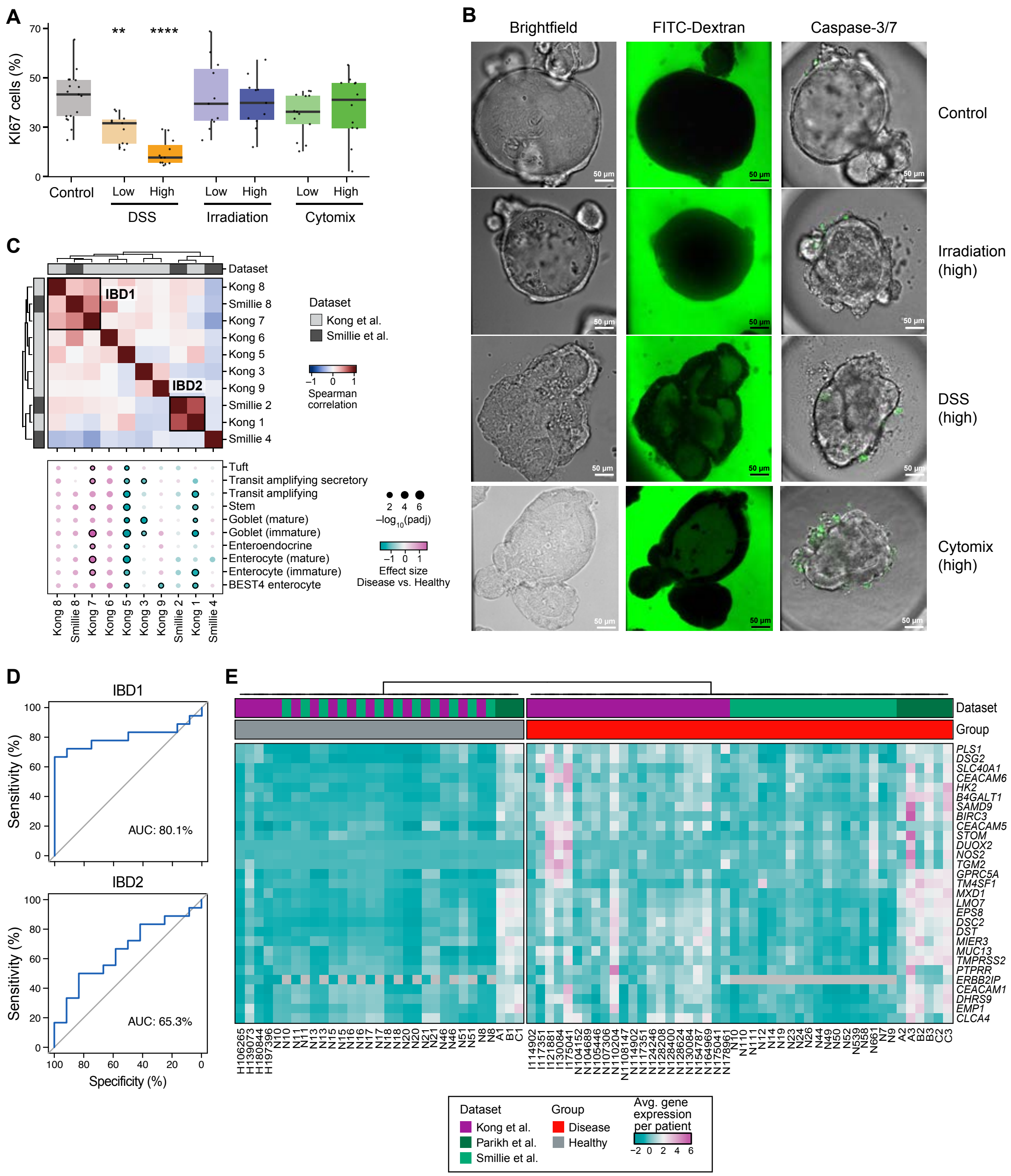


***Extended Data Figure 2: Comparison of organoid injury models to human IBD via functional analysis and derivation of IBD-relevant gene signatures.*** Related to Figure 1. **(A)** Quantification of proliferation one day post-injury, n>=10 organoids per condition. **(B)** Representative images of organoids used to quantify organoid permeability (4kDa FITC-dextran) and viability (activated Caspase 3/7) in Figure 1C. **(C)** Derivation of consensus gene programs from two IBD atlases [4,25]. Top: Spearman's correlation between GPs derived from IBD atlases, identifying GPs that are highly conserved between datasets, termed IBD1 and IBD2. Bottom: Effect size of GPs in diseased vs. healthy cells across epithelial cell types **(D)** Logistic regression model trained and validated on Ramnik et al. dataset using IBD1 and IBD genes to predict inflamed vs healthy. ROC represents hold out test on Smillie et al. **(E)** Average gene expression per patient for top 30 genes in GP IBD1 across healthy and diseased patients in three external datasets[4,25,46].

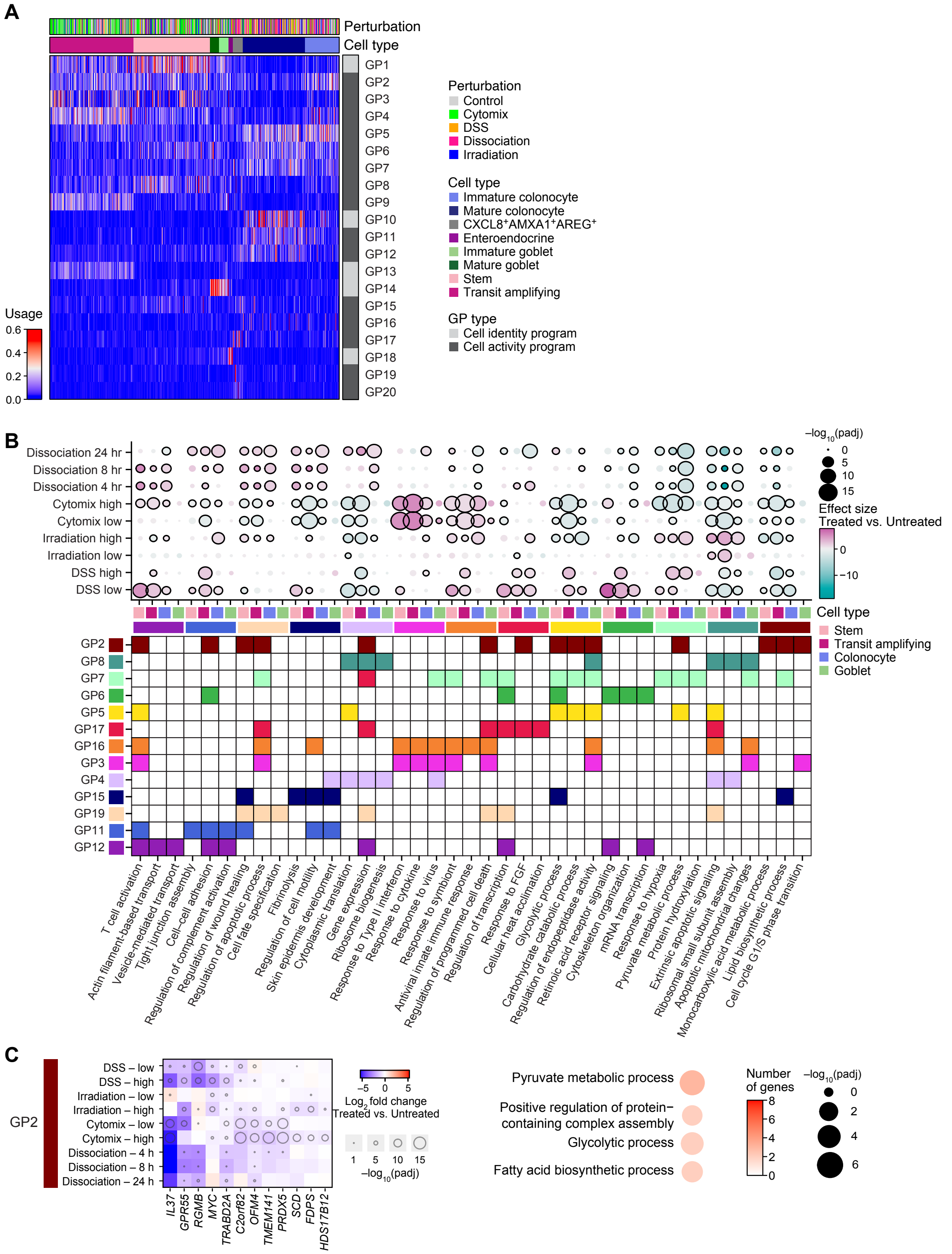


***Extended Data Figure 3: Identification of unbiased gene programs driving epithelial injuries in human colon organoids.*** Related to Figure 2. (**A**) Average usage of 20 gene programs identified through consensus NMF in each cell in the injured organoid atlas, clustered by cell type. GPs 1, 10, 13, 14, and 18 are classified as ‘cell identity’ rather than ‘activity’ programs based on their restriction to individual cell types and inclusion of canonical cell type-specific markers. **(B)** (Top) Effect size of cell activity GPs in each injury vs. control organoid, separated by cell type. Significant effect sizes (padj <= 0.05) are outlined in black. (Bottom) GO terms associated with each GP. **(C)** Left: Log2 fold change for genes in GP2 in each injury vs. control. Right: GO terms associated with genes in GP2.

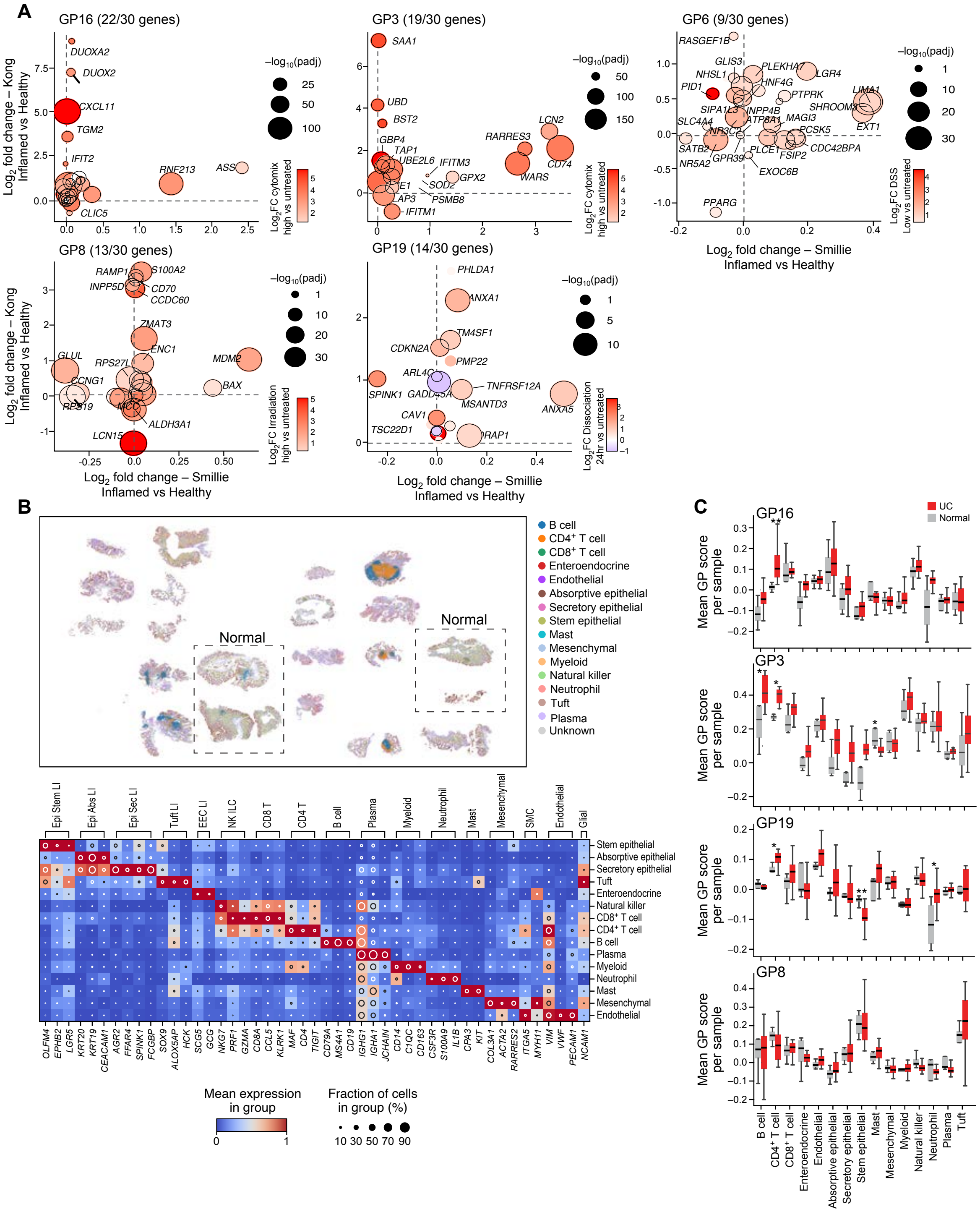

***Extended Data Figure 4: Relevance of injury-related gene programs to human inflammatory bowel disease.*** Related to Figure 2. **(A)** Logfold change of genes in injury-related GPs in transit amplifying cells from inflamed vs. healthy patients in the Smillie et al. dataset (x axis) vs. the Kong et al. dataset (y axis). Dots are colored by a gene's log fold change in injured vs. control organoids (GP16 = cytomix high, GP3 = cytomix high, GP19 = dissociation, 24 hr, GP6 = DSS low, GP8 = irradiation high). Black outline indicates padj <= 0.05 in organoids. Inset text depicts the number of genes within the top 30 genes per module with a positive log fold change (upregulated in inflamed vs. healthy) in both datasets. **(B)** Top: CosMx spatial tissue microarray of human colon samples from normal and ulcerative colitis patients. Cells are colored by epithelial cell type. Bottom: Expression of canonical cell type markers in the CosMx spatial tissue analysis. **(C)** Mean GP score per sample for injury-derived GPs 16, 3, 19, and 8, with n=4 normal samples and n=16 UC samples, *p <0.05, **p <0.01.

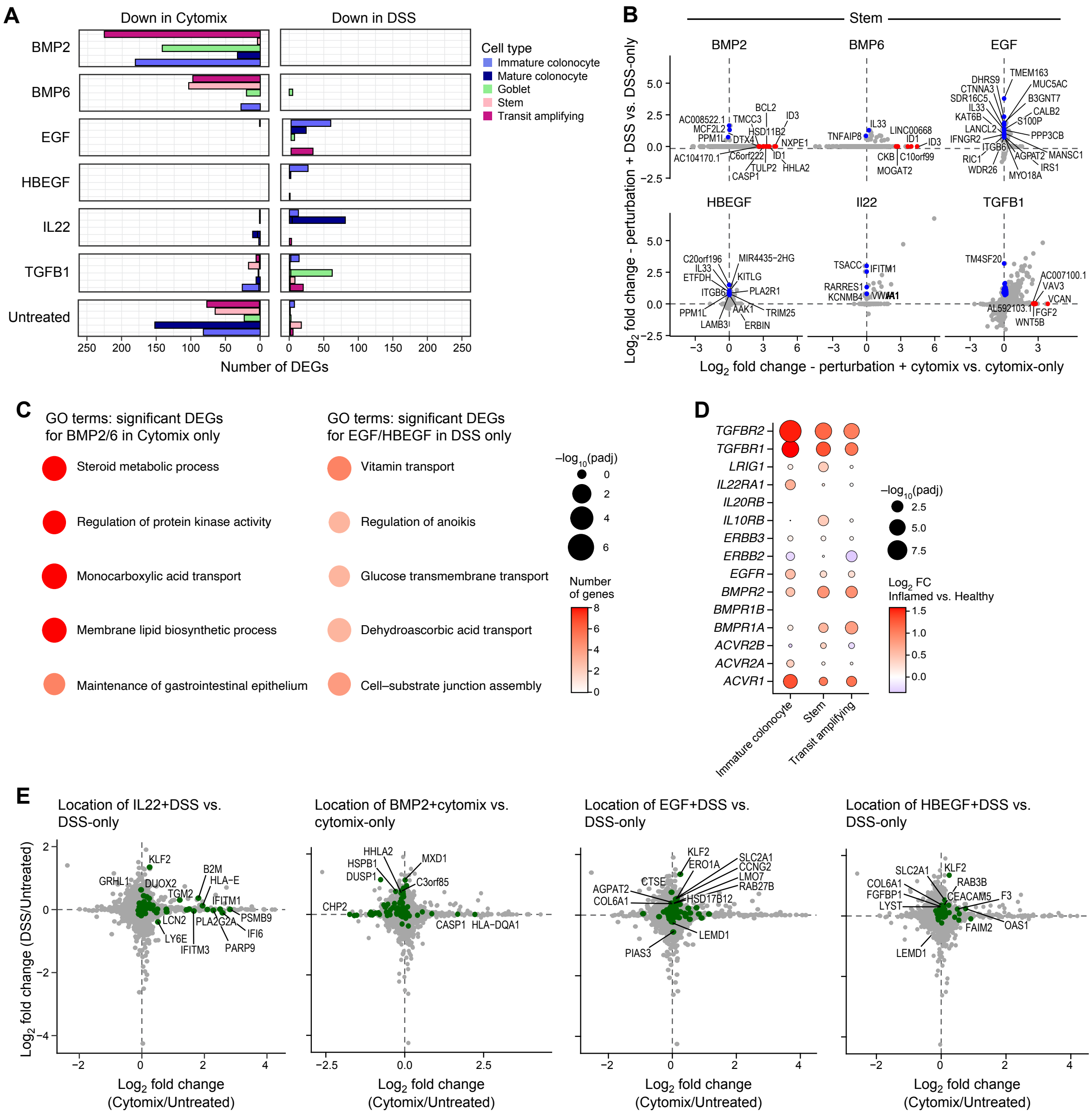

***Extended Data Figure 5: Comparison of organoid response to cytomix vs. DSS injury.*** Related to Figure 2. **(A)** Number of downregulated DEGs (p val <0.05, logFC > 1) for each secreted factor perturbation when applied to either the DSS model (right) or cytokine model (left). **(B)** Log2FC of DEGs in stem cells for each secreted factor applied to either the cytomix model (x axis) or the DSS model (y axis) (red=exclusively DE in cytomix-treated organoids, blue=exclusively DE in DSS-treated organoids). **(C)** GO terms associated with DEGs specifically upregulated in BMP2/6 in cytomix but not DSS-treated organoids, and in EGF/HBEGF in DSS but not cytomix-treated organoids. **(D)** Average log2-fold change of receptors and antagonists for applied perturbations in inflamed vs. healthy cells from human IBD tissue[25]. **(E)** Log2FC of DEGs in cytomix vs. untreated organoids (x-axis) vs. DSS vs. untreated organoids (y-axis). Green points depict DEGs upregulated by individual perturbations IL22, EGF, HBEGF in DSS-treated organoids, BMP2 in cytomix-treated organoids).

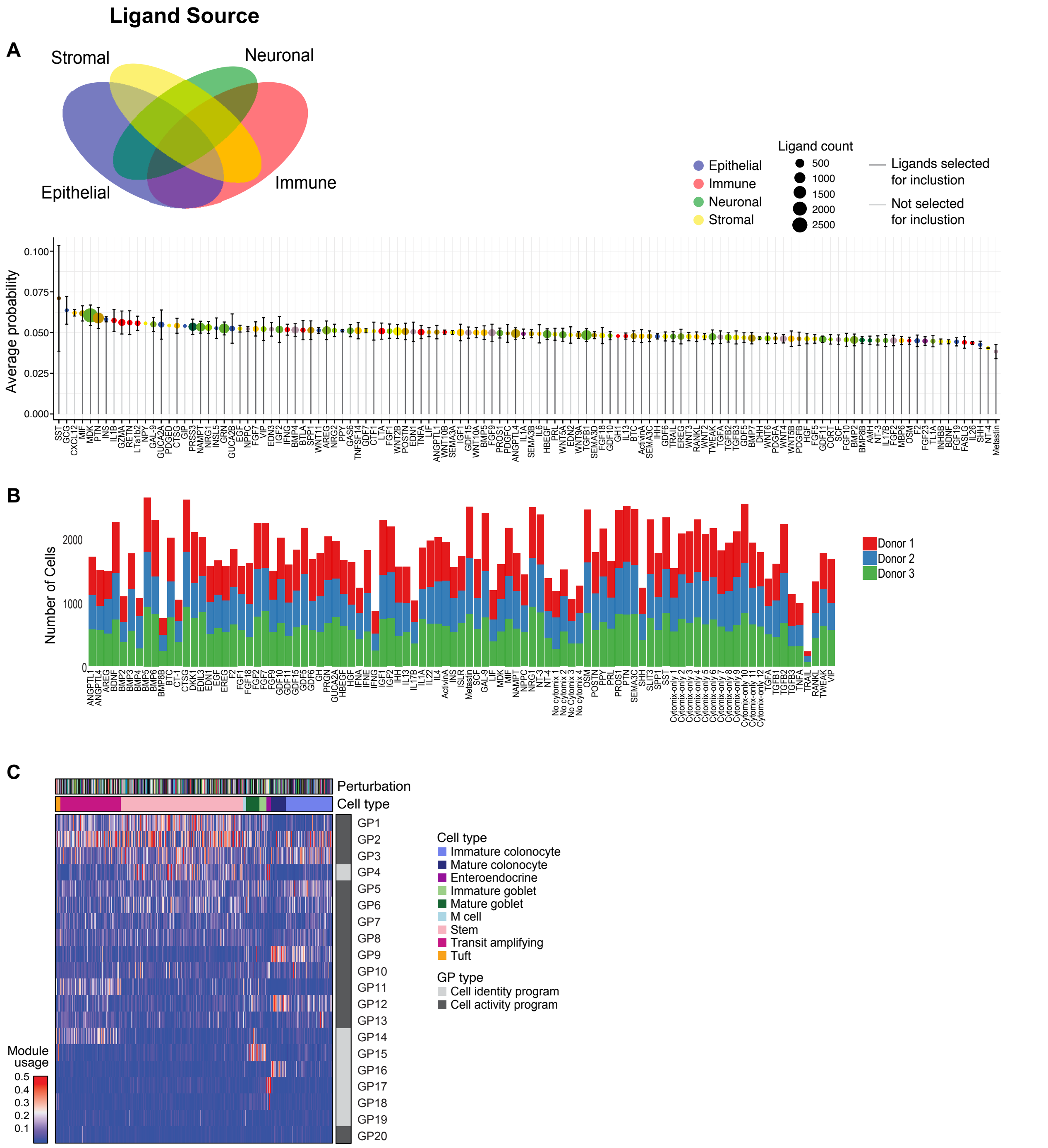


***Extended Data Figure 6: Ligand-receptor analysis prioritizes 79 secreted niche factors for screen.*** Related to Figure 3. **(A)** Average probability of interaction of niche secreted ligands with the epithelium during human colon during development[26,27], homeostasis[25,26], and disease[25]. Dots are colored based on the source of the predicted signal, and sized based on the total number of interactions predicted across 3 datasets. Ligands with dark gray stems were selected for inclusion in the screen. **(B)** Number of cells for each donor and perturbation in the final perturbation dataset, after filtering and QC. **(C)** Average usage of GPs in each cell in the perturbed organoid atlas, clustered by cell type. GPs are classified as ‘cell identity’ or ‘cell activity’ programs.

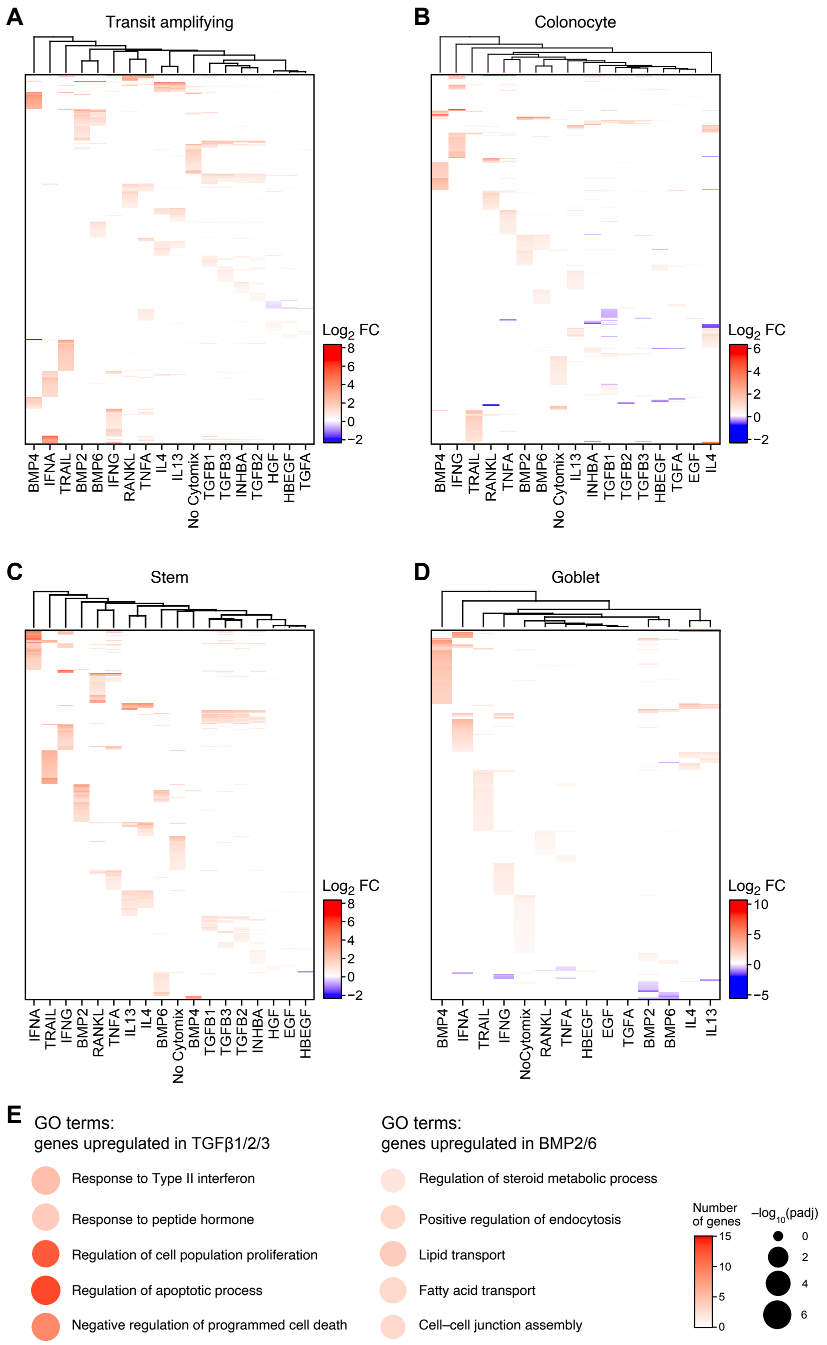


***Extended Data Figure 7: Differentially expressed genes associated with top perturbations reveal differences between similar ligand families.*** Related to Figure 3. **(A-D)** Log2FC of top 50 significantly upregulated DEGs in TA cells (A), colonocytes (B), stem cells (C) and goblet cells (D) for high impact perturbations (perturbation vs. cytomix-only control). **(B)** GO terms associated with DEGs upregulated across TGβ1/2/3 and BMP2/6 vs no-cytomix. DEGs were combined across colonocytes and stem cells.

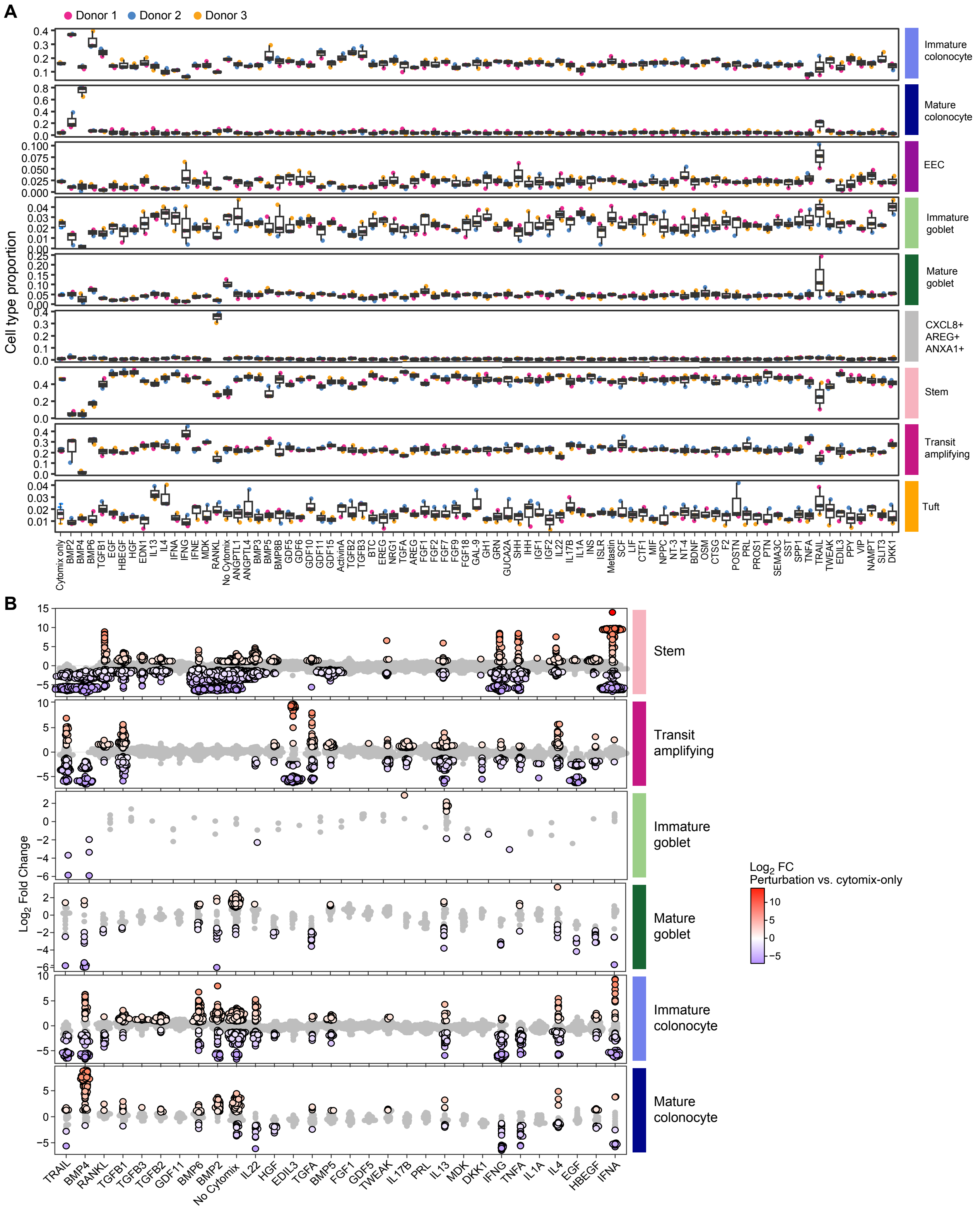


***Extended Data Figure 8: Subset of perturbations leads to significant changes in cell type composition.*** Related to Figure 4. **(A)** Cell-type composition in perturbation organoid dataset on a per donor basis. **(B)** Log fold changes of neighborhoods within each cell type for each perturbation vs no-cytomix, calculated via differential abundance testing (using MILO). Outlined points denote significance (FDR <= 0.05).

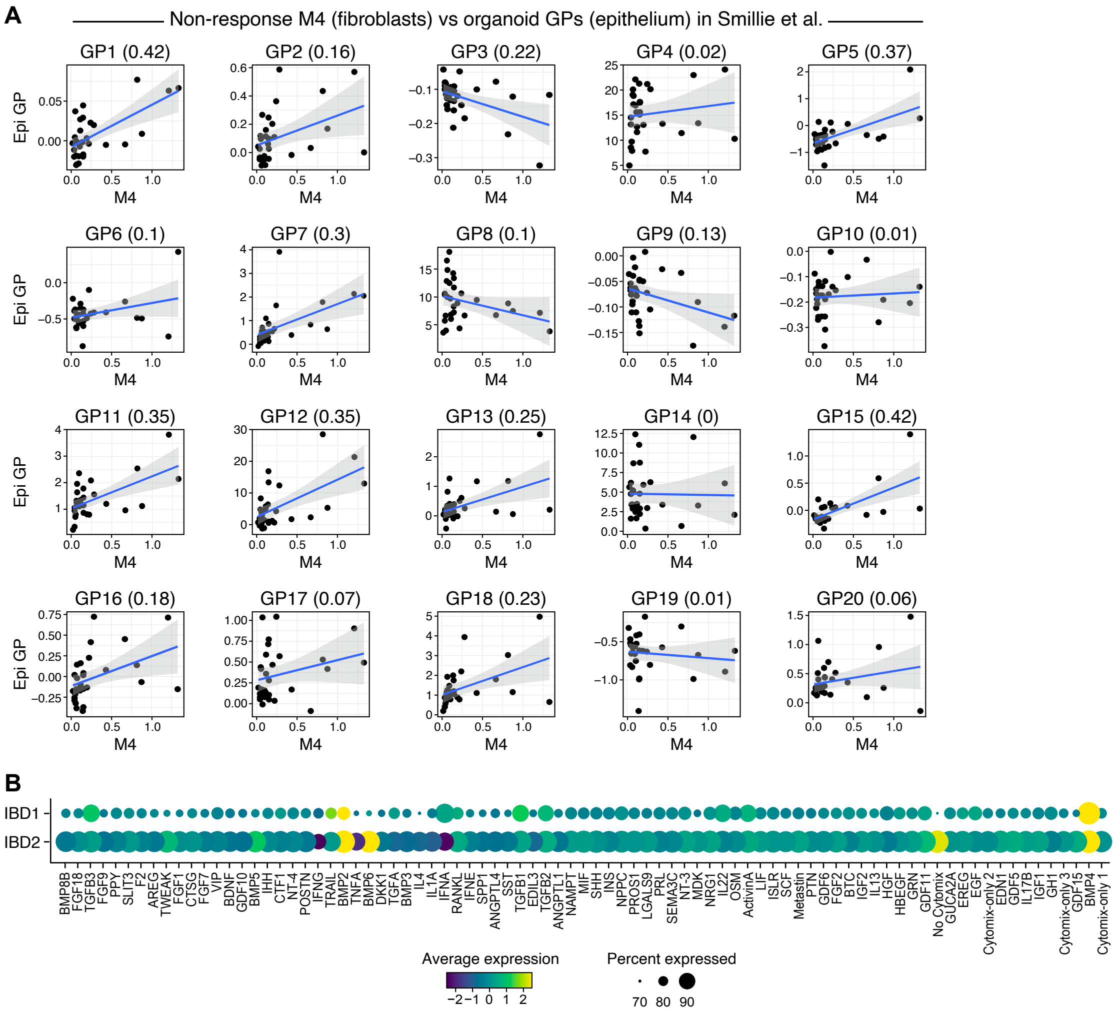


***Extended Data Figure 9: Assessment of IBD-relevant gene programs within treated organoids.*** Related to Figure 5. **(A)** Average expression per patient of an anti-TNFα non-response signature in fibroblasts[54] (M4, x-axis) vs. the average expression per patient of organoid-derived GPs in the epithelium (y-axis), in the Smilie et al. atlas[4]. $R^2$ values indicated in brackets. **(B)** Scaled expression of GPs IBD1 and IBD2 (Fig. 1) in each organoid perturbation.
